# A broad-spectrum macrocyclic peptide inhibitor of the SARS-CoV-2 spike protein

**DOI:** 10.1101/2022.11.11.516114

**Authors:** Vito Thijssen, Daniel L. Hurdiss, Oliver J. Debski-Antoniak, Matthew A. Spence, Charlotte Franck, Alexander Norman, Anupriya Aggarwal, Nadia J. Mokiem, David A. A. van Dongen, Stein W. Vermeir, Minglong Liu, Wentao Li, Marianthi Chatziandreou, Tim Donselaar, Wenjuan Du, Ieva Drulyte, Berend-Jan Bosch, Joost Snijder, Stuart Turville, Richard J. Payne, Colin J. Jackson, Frank J. M. van Kuppeveld, Seino A. K. Jongkees

## Abstract

The ongoing COVID-19 pandemic has had great societal and health consequences. Despite the availability of vaccines, infection rates remain high due to immune evasive Omicron sublineages. Broad-spectrum antivirals are needed to safeguard against emerging variants and future pandemics. We used mRNA display under a reprogrammed genetic code to find a spike-targeting macrocyclic peptide that inhibits SARS-CoV-2 Wuhan strain infection and pseudoviruses containing spike proteins of SARS-CoV-2 variants or related sarbecoviruses. Structural and bioinformatic analyses reveal a conserved binding pocket between the receptor binding domain, N-terminal domain and S2 region, distal to the ACE2 receptor-interaction site. Our data reveal a hitherto unexplored site of vulnerability in sarbecoviruses that peptides and potentially other drug-like molecules can target.

**Significance statement:** This study reports on the discovery of a macrocyclic peptide that is able to inhibit SARS-CoV-2 infection by exploiting a new vulnerable site in the spike glycoprotein. This region is highly conserved across SARS-CoV-2 variants and the subgenus sarbecovirus. Due to the inaccessability and mutational contraint of this site, it is anticipated to be resistant to the development of resistance through antibody selective pressure. In addition to the discovery of a new molecule for development of potential new peptide or biomolecule therapeutics, the discovery of this broadly active conserved site can also stimulate a new direction of drug development, which together may prevent future outbreaks of related viruses.

## Introduction

As of February 2023 severe acute respiratory syndrome coronavirus 2 (SARS-CoV-2) has infected more than 755 million individuals and caused over six million deaths (https://covid19.who.int/). Despite the rapid development and deployment of vaccines, antigenic evolution of SARS-CoV-2 has led to a resurgence in COVID-19 cases globally. As such, effective prophylactic and treatment options for COVID-19 are still needed to supplement vaccination efforts and ensure preparedness for future coronavirus outbreaks, including future variants against which current vaccines may be less effective. Currently approved small molecule antivirals include remdesivir^1^ and molnupiravir,^2^ both of which target the RNA-dependent RNA polymerase, and nirmatrelvir,^3^ a covalent inhibitor of the SARS-CoV-2 main protease. Importantly, both targets are only expressed when the virus has already infected the cell, thus these molecules cannot prevent initial infection. An alternative antiviral strategy is through the use of monoclonal antibodies that block early events in the SARS-CoV-2 life cycle by targeting the trimeric spike (S) glycoprotein,^4^ a class I fusion protein that can be broadly divided into two subunits: S1 and S2.^5^ The S1 subunit comprises two domains, S1A and S1B, the latter of which binds to the host angiotensin-converting enzyme 2 (ACE2) receptor.^6,7^ Subsequently, the S2 subunit undergoes large-scale conformational changes to drive fusion of the viral and host membranes and initiate an infection.^8^ The most potent monoclonal antibodies in clinical development target the S1B domain and interfere with ACE2 binding.^9^ However, these antibodies are particularly susceptible to mutations found in SARS-CoV-2 variants of concern (VOCs). For example, the unprecedented number of mutations found in the Omicron BA.1 and BA.2 S1B domain have severely impacted the efficacy of six out of nine clinical-stage monoclonal antibodies^10^. The least affected antibody, Sotrovimab,^11^ was found to have a 3-fold loss in potency against Omicron BA.1 and 27-fold loss against BA.2 whereas two other antibodies, Cilgavimab^12^ and Adintrevimab,^13^ showed a 20-fold decrease in activity.

Additionally, Omicron sublineages BA.2.12.1, BA.4/5 and BA.2.75 display enhanced transmission and immune evasion compared to BA.2.^14,15^ Antibodies are also expensive to produce at scale and have strict cold-chain requirements that lead to high treatment costs. This highlights the difficulty in antibody development against mutating viruses that are under selection pressure from the immune system and the need to develop broad-spectrum prophylactics and therapeutics using alternative molecular modalities.

Macrocyclic peptides are a class of molecules that retain much of the binding affinity and selectivity of antibodies (or single-chain antibodies such as nanobodies) in a much smaller molecule (ca. 2-3 kDa vs ∼150 kDa for monoclonal antibodies and 15 kDa for nanobodies). Such smaller biomolecules can access binding sites that larger molecules cannot, and also possess superior tissue penetration.^16^ Importantly, high-affinity cyclic peptides can be generated with attractive drug-like parameters, such as protease stability, through the use of peptide display technologies (e.g. phage and mRNA display) with genetic code reprogramming and/or chemical modification on the peptide libraries.^17^ In this work we report the use of mRNA display under a reprogrammed genetic code for the discovery of a 17 amino acid macrocyclic peptide that binds with high affinity to a conserved region of the spike protein. This macrocyclic peptide neutralizes SARS-CoV-2 independently of ACE2-binding inhibition by targeting a previously undescribed druggable site, and can prevent infection across all tested SARS-CoV-2 VOCs and other sarbecoviruses with little to no loss of activity.

## Results and Discussion

We used the RaPID (Random non-standard Peptide Integrated Discovery) system^18^ to assemble two peptide libraries containing approximately 10^12^ different sequences. These peptides are tagged during translation with their encoding mRNA and are cyclized by recoding of methionine to *N*-chloroacetylated D-tyrosine or L-tyrosine (one for each library), which cyclizes with a cysteine residue placed after 15 randomized codons that encode any of the 19 remaining L-amino acids (excluding methionine). Panning of these libraries with prefusion-stabilized WT SARS-CoV-2 S ectodomains (Wuhan-Hu-1) immobilized on magnetic beads recovered binding peptides, and the cDNA of these was amplified to provide input mRNA for a subsequent round. Following five iterative rounds of enrichment, fetal calf serum was added prior to the incubation with the spike for a further two rounds to improve serum stability (Fig. 1A and S1-4, Table S1). In a parallel approach, the input of the third round (where enrichment was first observed) was panned with the spike S1B domain, which contains the ACE2 receptor-binding site. High throughput next-generation sequencing of all enriched libraries revealed sequences that were then tallied, aligned, and clustered by sequence similarity to select potential cyclic peptide ligands (Fig S5), 15 of which were produced by solid-phase peptide synthesis (Table 2, Fig S6). Purified synthetic cyclic peptides were then tested directly for activity in a pseudovirus assay for the ancestral SARS-CoV-2 Wuhan-Hu-1 strain (Fig. 1B and S7). Two of the tested sequences (S1b3inL1 and SARS2L1, both from the L-tyrosine initiated library), showed pseudoviral neutralization activity at sub-micromolar concentrations. These peptides were selected for further study, along with a close homologue of S1b3inL1 (coded S1b3inL4). A protein thermal shift assay using either trimeric spike ectodomains (Wuhan-Hu-1) or the S1B-domain alone confirmed that S1b3inL1 binds to the receptor binding domain (Fig. S8). For SARS2L1, a thermal shift was only observed for the full trimeric spike, indicating that it binds to a region outside of the S1B or recognizes a quaternary epitope. Several other non-inhibitory peptides also showed a thermal shift, indicating that these are binding to non-inhibitory regions (Fig. S9 and Table S3). A competitive enzyme-linked immunosorbant assay (ELISA) showed that neither active peptide prevented spike binding to ACE2, which suggests both might be binding to an auxiliary site (Fig. 1C). We also tested for inhibition of genuine virus infection using the ancestral Wuhan strain by measuring cytopathic effects in HEK293 ACE2+ TMPRSS2+ cells (Fig. 1D). The peptide S1b3inL1 (Fig. 1E) showed good activity with an *EC*_*50*_ of 5.8 µM, with the homologous S1b3inL4 exhibiting less potent viral neutralization (11.4 µM). We did not observe detrimental off-target effects with the S1b3inL1 peptide, rather we observed some unexplained stimulation of cell growth at the highest concentrations, and subsequent toxicity assays showed only modest toxicity (Fig. S10-11, detectable at concentrations above 50 μM). While SARS2L1 clearly showed inhibition of viral entry, the *EC*_*50*_ curve did not reach a plateau due to solubility limitations. In past experiments we have seen a good correlation between trends with whole virus and pseudotyped virus data,^19^ and so with this activity validated further experiments were carried out using the safer pseudotyped viruses. Based on these data, we focused our attention on understanding the potentially unique mechanism of action of the most promising cyclic peptide S1b3inL1.

**Fig. 1.**
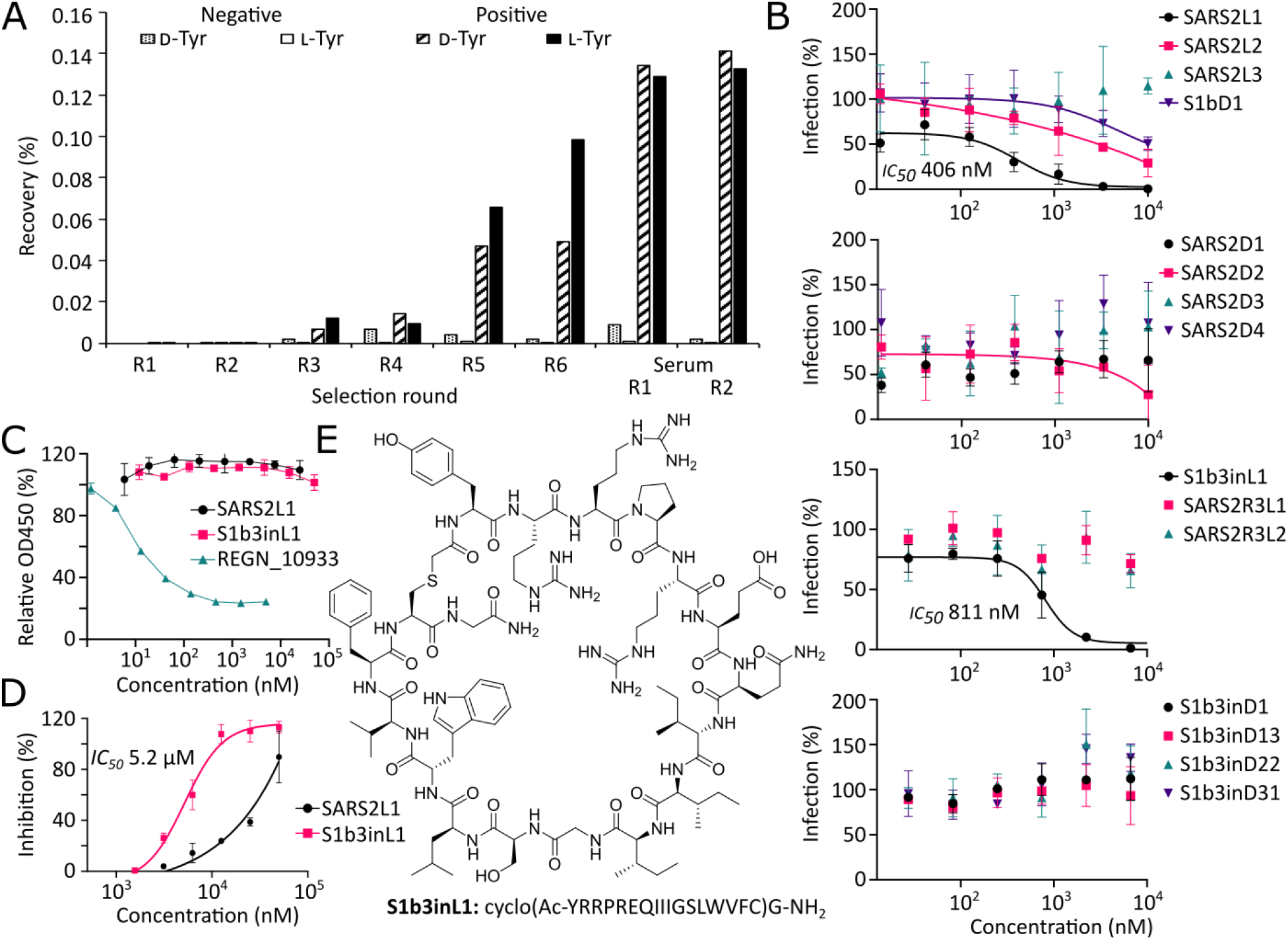
Macrocyclic peptide selection strategy and selection results. A) Enrichment of library recovery across rounds for two random 15-mer peptide libraries initiated with chloroacetyl-D-tyrosine and -L-tyrosine. Rounds with fetal calf serum added during incubation are marked as ‘serum’. B) Dose-response curves from screening of peptides identified by high throughput sequencing of enriched libraries for inhibition of pseudovirus infection in VeroE6 cells. C) ELISA showing lack of competition for ACE2 binding to spike protein by the most promising peptide hits, compared to REGN_10933 antibody positive control. D) Dose-response curve for SARS-CoV-2 inhibition assays in HEK293 ACE2+ TMPRSS2+ cells with the most promising hits from the pseudovirus screen. E) Structure of most promising hit, peptide S1b3inL1, and its sequence.

To understand how the peptide S1b3inL1 interacts with the SARS-CoV-2 spike glycoprotein, we used hydrogen-deuterium exchange (HDX) footprinting on the S1 protomer (Fig. S12). This revealed a putative binding site formed by residues 388-399, 421-431 and 511-520, which localize to a region of the S1B that is distal to the ACE2 interacting region. To obtain a detailed understanding of the binding site and macrocyclic peptide structure, we performed cryo-electron microscopy (cryo-EM) single-particle analysis on 6P-stabilized spike ectodomains incubated with S1b3inL1. Three-dimensional classification revealed a single, well-defined class consisting of the prefusion spike with two S1B domains in the open conformation and one in the closed conformation (Fig. S13). Subsequent processing steps yielded a reconstruction with a global resolution of 3.2 Å, which showed additional density consistent with the size of the macrocyclic peptide on each of the three S1B domains. The two peptide-bound S1B domains in the open conformation were weakly resolved in the cryo-EM map due to conformational variability. As such, a focused classification was performed on the better resolved closed S1B domain, which improved the local resolution to ∼3.5 Å, sufficient to model the entire cyclic peptide and surrounding spike residues (Figs. 2A-B, S14 and Table S4). The bioactive conformation of S1b3inL1 is stabilized by hydrophobic sidechain packing in the peptide core and hydrogen bonding between the sidechain of R5 and the backbone carbonyl groups of I8 and G11 (Fig. 2C). The cyclic peptide interacts with a recessed region of the S1B domain through shape complementarity, burying 744 Å^2^ of its solvent-accessible surface area (∼38% of total). Consistent with our HDX-MS analysis, the convex face of the cyclic peptide is juxtaposed with residues 353, 355, 390-392, 426-430, 462-466 and 514-519. The binding of S1b3inL1 to the S1B domain is primarily mediated by hydrophobic interactions and stabilized by backbone hydrogen bonding to S1B domain residues R466, F456 and L517. When the S1B domain is in the closed conformation, the peptide is brought into proximity with the N-terminal domain (S1A) and S2 residues from the adjacent spike protomers (Fig. 2D). Specifically, S1b3inL1 interfaces with residues 39-40, 51-53, 197-198, 751, 755, 969, 973-974, 979, 983 and the N234 glycan (Figure 2D). Collectively, this buries an additional 260 Å^2^ of the solvent-accessible surface area of the cyclic peptide (∼52% in total). We also observed putative solvent densities within the quaternary binding site which could conceivably mediate hydrogen bonding between the peptide and surrounding spike residues. However, we opted to leave these unmodelled due to the limited resolution. Notably, the part of the binding site for S1b3inL1 found in the S1B domain has substantial overlap with the pan-sarbecovirus ‘site V’-binding antibody S2H97.^20^ However, unlike S1b3inL1, S2H97 requires extensive opening of the S1B domain to expose this cryptic epitope. As such, the ternary S1B3inL1 binding site formed by S1A, S1B and S2 represents a hitherto unexplored site of vulnerability.

**Fig. 2.**
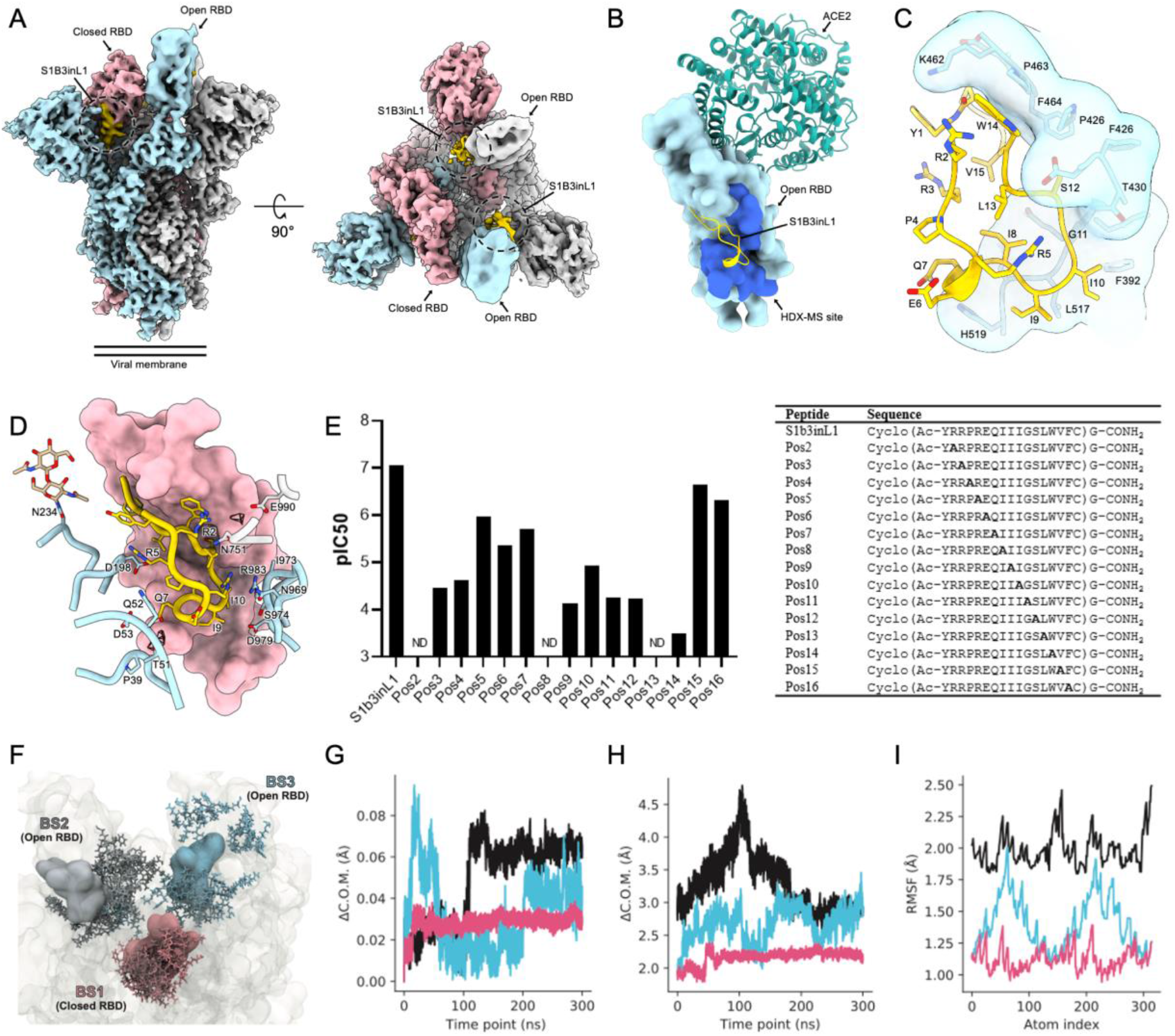
Characterization of the S1B3inL1 binding site. A) Composite cryo-EM density map for the SARS-CoV-2 spike ectodomain in complex with S1B3inL1, shown as two orthogonal views. The spike protomers are colored blue, gray, and pink, and the macrocyclic peptide is colored gold. (B) Surface representation of the S1B3inL1-bound S1B overlaid with the S1B-bound ACE2 (PDB ID: 6M0J). Residues 388-399, 421-431 and 511-520, identified by HDX-MS, are colored dark blue. C) Close-up view showing the interactions between S1B3inL1 and the SARS-CoV-2 S1B. D) Close-up view of the S1B3inL1-bound closed S1B. Selected S1A and S2 residues are shown as sticks, and putative solvent densities are shown as a mesh. E) Relative SARS-CoV-2 pseudovirus inhibition activity of peptides with residues at positions 2 to 16 substituted for alanine, with sequences of peptides used in the assay displayed in the table. Activities are shown as pIC_50_ values, with a higher pIC_50_ value indicating more potent activity. Alanine mutations for which no inhibition was detected are denoted as ‘ND’. F) Initial poses for S1b3inL1 in binding sites (BS) 1-3 (colored by spike protomer as in panel A) are shown in surface representation. Coordinates for final poses after 300 ns are shown in licorice representation. Initial coordinates of the full-length spike are shown in transparent surface representation. G) Center of mass (COM) evolution for S1B3inL1 BS1-3 from a single representative trajectory. BS2 and 3 peptides (cyan and black) exhibit large COM movements over the 300 ns trajectory, relative to BS1 peptide. H) Distance between the COM of S1B3inL1 in BS1-3 and the COM of the closest S1B in the initial coordinates. S1B3inL1 COM movements in BS2 and 3 are decoupled from COM movements in the S1B. BS1 remains stable with the closed S1B. I). Root mean square fluctuation (RMSF) over all S1B3inL1 atoms averaged over each sampled trajectory. S1B3inL1 in BS1 remains more conformationally stable than BS2 and 3.

To further understand which of these interacting amino acids are important for the antiviral activity of S1b3inL1, we performed an alanine scan on all positions varied in the library (residues 2-16) and used this to evaluate the contribution of each amino acid to activity in a pseudovirus neutralization assay (Fig. 2E). Positions 15 and 16, which are solvent exposed in both the open and closed conformation, showed high tolerance for substitution with alanine. A modest decrease in antiviral activity was observed for positions 5, 6 and 7, which do not directly interact with the S1B domain but may interact with S1A or S2 residues in the closed conformation (with position 5 being involved in stabilizing the conformation of the cyclic peptide). All other positions show substantial or even total loss of activity when amino acids were mutated to an alanine. Mutation at positions 2, 8 and 13 led to a complete loss of detectable activity; of these, only residue 8 is positioned at the S1B domain-interacting interface and thus mutation here probably leads to loss of binding for both the open and closed state. The sidechain of L13 is oriented towards the peptide core, so mutation at this site may compromise the bioactive conformation of S1b3inL1. The sidechain of R2 is juxtaposed with W14 and may stabilize it in a conformation favorable to S1B domain binding. However, R2 may also participate in solvent-mediated interactions with S2 residue E990, and thus the mutation at this position could affect the binding of the buried “down” state. Collectively, the alanine scan data suggests that the buried binding mode contributes to the antiviral mechanism of the peptide.

The hypothesized mechanism of action of S1b3inL1 was further supported by 6 independent replicates of 300 ns (3 µs total sampling time) molecular dynamics simulations carried out on the whole glycosylated spike protein, with peptide bound to all three binding sites (Fig. 2F and S15). This showed that the peptides bound to the “up” states are more prone to drift out of the modelled bindings site, while those bound to the “down” state are much more stable (Figs. 2G-I). Based on these observations, we hypothesize that S1b3inL1 exerts its antiviral activity by stabilizing the “down” conformation through the buried binding mode and thereby modulating the conformational landscape of the spike protein. Conceivably, by targeting a quaternary epitope and stabilizing one S1B domain in a closed conformation, S1b3inL1 could interfere with the essential conformational changes that are critical for viral membrane fusion.^21^

The identified binding site for S1b3inL1 in the spike protein is buried and distant from residues that are mutated in all currently identified VOCs (Fig. 3A), suggesting that this site might be less susceptible to antigenic drift. To investigate this, we performed phylogenetic analysis of coronavirus spike proteins and sequence entropy analysis (Fig. 3B and S16-17). The data show that the binding site is well conserved across SARS-CoV-2 VOCs. This conservation also extends to sarbecoviruses and norbecoviruses, but not all betacoronaviruses. This is thus a new functional region of the S protein that may be broadly exploitable within these related families through a similar approach. Indeed, surface plasmon resonance (SPR) analysis showed that mutations in several S proteins in VOCs had negligible influence on the binding of S1b3inL1 (Fig. 3C and S18). Peptide S1b3inL1 was able to bind to the spike with *K*_*d*_ around 50 nM, while the less active peptide S1b3inL4 showed correspondingly lower affinity binding (*K*_*d*_ around 200 nM). These two peptides differ in only a few residues, most of which are conservative changes (Q7L, I8V, L13V, V15N). The two non-conservative changes (R3V and E6K), however, both point away from the S1B interaction surface. Given that the alanine scan also shows a poor tolerance for alanine substitutions at other residues outside the S1B binding interface (especially R2 and R3), and that the MD simulations show that the peptides bound to the open S1B domain tend to dissociate, these results together indicate that the binding seen in the thermal shift assay to the S1B domain alone is not sufficient to explain the activity. The SARS2L1 peptide was also found to bind exceptionally strongly (0.8 nM). Why this strong binding and high activity in pseudovirus assays does not translate to equally potent neutralization of genuine virus infection remains unknown, and is the subject of ongoing investigation. The high conservation within the binding site we identify here correlates with broad activity of S1b3inL1 in neutralization assays with pseudotyped viruses of several VOCs (Fig. 3D). The *EC*_*50*_ values were seen to fluctuate slightly, but activity was retained against Alpha (B1.1.1.7; 1.5 fold), Beta (B.1.351; 3.1 fold), Delta (B.1.617.2; 2.1 fold) and even the heavily mutated Omicron (B.1.1.529; 3.1 fold for BA.1 and 2.1 fold for BA.2) variants. This was also seen to extend beyond SARS-CoV-2 (Fig. 3E), with infection inhibition being largely unchanged with pseudoviruses displaying spike proteins from other sarbecoviruses (SARS1-S, 1.3 fold, and WIV-16-S, 4.1 fold). Taken together, our data show that cyclic peptide S1b3inL1 has exceptionally broad activity for such a small spike-binding molecule and suggests that this auxiliary binding site on the S protein might serve as a privileged target for drug discovery efforts for future pandemic outbreaks of coronaviruses.

**Fig. 3.**
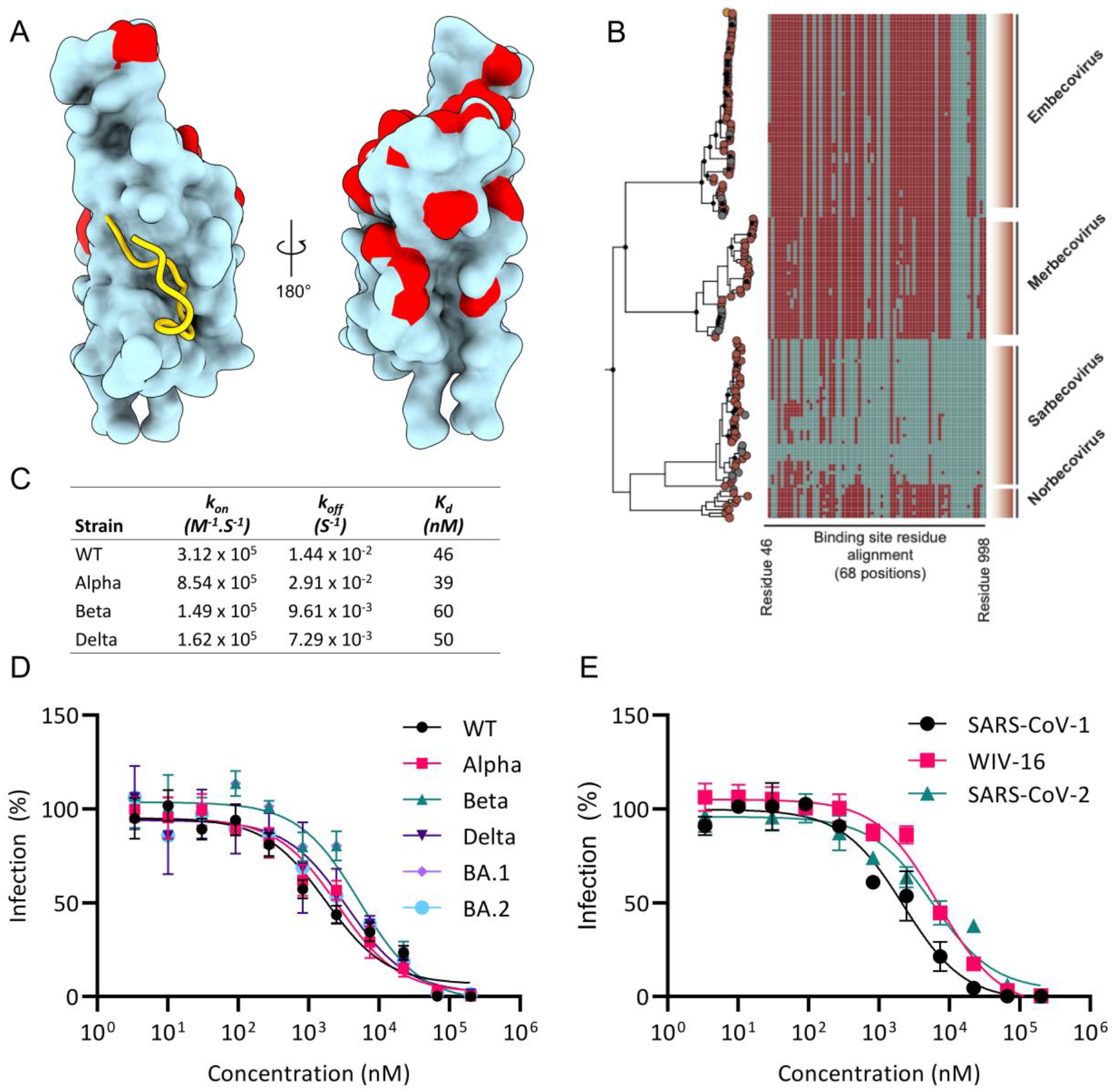
Sequence conservation of the S1B3inL1 binding site. A) Close-up view of the S1b3inL1 binding site (gold cartoon) relative to mutation sites (red) in the Alpha, Beta, Gamma, Delta, Lambda, Mu and Omicron (BA.1, BA.2, BA.2.12.1, BA.2.75, and BA.4/5) variants within the S1b domain (cyan surface representation). B) Maximum likelihood phylogenetic reconstruction of β-coronavirus lineages. The topology is rooted at the midpoint and is reconstructed from 725 non-redundant amino acid sequences. Peptide binding site residues (designated as SARS-COV2 residue, cyan, or non-SARS-COV2 residue, maroon) are aligned respective tips. Major β-coronavirus lineages are labeled. C) Characterization of S1b3inL1 binding to S1B by Biacore for wild type and variants. D) Dose-response curve for inhibition of pseudovirus infection in VeroE6 cells by S1b3inL1 across VOCs. E) Dose-response curve for inhibition of pseudovirus infection in VeroE6 cells by S1b3inL1 across sarbecoviruses.

While cyclic peptide S1b3inL1 shows broad activity, improving its potency to the point where it could be used in the clinic remains a challenge. As a first step towards this, S1b3inL1 was dimerized with a varying number (8 or 16 total) of a polyethylene glycol-based amino acid spacer connecting two peptides at their *C*-termini (2-[2-[2-(amino)ethoxy]ethoxy]acetic acid, ‘AEEA’, denoted AEEA8 and AEEA16, Fig. 4A, S18-20, and Table S5). We considered that dimerization might increase antiviral potency in several possible ways, such as by bridging two binding sites within the same or between adjacent S proteins, or by decreasing the off-rate through increasing the likelihood of re-binding. The number of spacers was based on a rough approximation of the distance between two binding sites and accounting for the flexibility of the spacer, with the two different lengths potentially allowing access to different ways of improving binding. Dimerization resulted in a dramatic increase in antiviral activity, i.e. up to 2 orders of magnitude (Fig. 4B-C), but the length of the linker had little influence on activity and it thus seems unlikely that these are bridging two binding sites as this would be expected to have a clear length optimum. The best of these dimers inhibited the infection by WT pseudovirus with an *EC*_*50*_ of 0.01 µM (10 nM, 240-fold improvement with BA.2), and activity again remained consistent across the other sarbecoviruses tested. We therefore believe this to be a promising strategy to improve the potential therapeutic window of this peptide, but these molecules were found to be substantially lower in solubility than the macrocycle alone at physiological pH (despite the use of a flexible polar linker) and this has precluded animal studies. Additional linker and/or peptide sequence optimization will likely be needed for testing to proceed further, but this molecule has clearly demonstrated its promise and connection to peptides binding at another site may represent a means to further enhance antiviral activity.

**Fig. 4.**
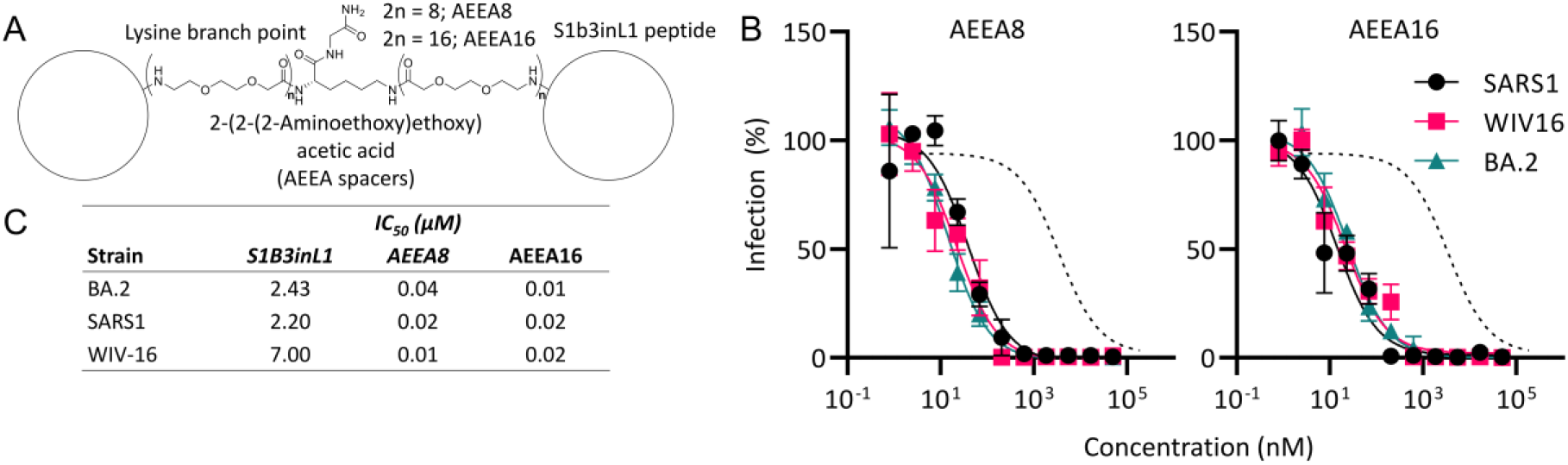
Dimerization for improved potency. A) Representative structure of S1b3inL1 AEEA dimers. B) Dose-response curve for inhibition of pseudovirus infection in VeroE6 cells by S1b3inL1 dimers of varying length (total number of AEEA spacers as indicated). Dotted lines indicate fit to monomeric S1b3inL1 peptide with BA.2 (Fig. 3D) C) *IC*_*50*_ values for S1b3inL1 dimers in the pseudovirus infection assay in panel B.

## Conclusions

We report here the discovery of a macrocyclic peptide that can inhibit infection across an broad range of SARS-CoV-2 variants by exploiting a previously unknown vulnerability that has high sequence conservation. Other approaches have been reported for developing spike-binding molecules other than antibodies,^22^ but these are still much bigger than the peptide reported here. Nanobodies are much smaller than antibodies but are still derived from them and typically discovered through the same immunization approach and so are unlikely to bind to significantly different epitopes. While potent nanobodies for the spike have been found that bind multiple different epitopes on the S trimer,^23–30^ none of these appear to bind to the same site as S1b3inL1 and those that partially overlap appear to only bind the S1B domain when it is in the ‘up’ conformation.^23^ The multivalent DARPin ensovibep^31^ can resist mutational changes through cooperative binding of three S1B-binding modules with subtly different paratopes to the ACE2 binding site. However, the immune privileged site found in the present study gives even broader activity and may not be accessible to even the smallest antibody mimetics. Aptamers against the spike protein have also been discovered,^32–34^ but these are again very large molecules that likely cannot access binding sites different from those of antibodies and are oligonucleotides are difficult to translate into therapeutics.^35^ Finally, miniproteins designed to bind the S1B domain give very potent inhibition of infection,^36^ but because these were designed to interact with the ACE2 binding site they are susceptible to mutations which frequently arise here. Small macrocyclic peptides thus appear to have key advantages over these competing biological approaches in this setting. A protein design approach to either improve the existing S1b3inL1 peptide or find new molecules that can exploit the new vulnerable ternary binding site elucidated in our work may prove to be a powerful strategy for the development of *bona fide* drug candidates in the future. Other peptide discovery campaigns from random peptide libraries, such as by mRNA display,^37^ affinity selection – mass spectrometry^38^ or covalent phage display,^39^ have found peptides that can bind to the spike protein S1B domain, but these have not resulted in bioactive sequences. We consider that our combination of a more sophisticated selection strategy using full spike protein followed by sub-domains, together with cluster-based hit identification following high-throughput data analysis and functional follow-up screening, as potential reasons for the successful identification of cyclic peptides with antiviral activity in this work.

The broadly active macrocyclic peptide S1b3inL1 described here, as well as the new druggable site it reveals, demonstrates the power of *in vitro* selection technologies to provide alternative solutions to those offered by small molecules or biologicals. Our high-resolution peptide-bound cryo-EM structure of S1b3inL1 gives a valuable perspective on the binding of the novel antiviral cyclic peptide to the full trimeric S protein target. Further mutagenesis and computational studies on this binding site show the quaternary interaction to be the likely active form, and bioinformatic analysis shows that this site is well conserved. We posit that the wide ternary binding pocket exploited here would be difficult to exploit with a traditional small molecule drug and note that no reported antibody or nanobody has been able to enter this occluded site despite several binding adjacent to it. Our work has revealed a potentially important druggable site in the coronavirus S protein that can spur development of new treatment approaches, as well as a molecule that can exploit this for all known variants of SARS-CoV-2 and several other sarbecoviruses, and so represents a valuable contribution to future pandemic preparedness.

## Acknowledgments

We thank J. A. W. Kruijtzer for assistance with peptide synthesis, and the Utrecht Sequencing Facility for providing sequencing service and data. We thank C. A. M. de Haan, W. Li and Y. Lang for advice and technical assistance.

## Funding

This work was partially funded by the Corona Accelerated R&D in Europe (CARE) project. The CARE project has received funding from the Innovative Medicines Initiative 2 Joint Undertaking (JU) under grant agreement No 101005077. The JU receives support from the European Union’s Horizon 2020 Research and Innovation Programme, the European Federation of Pharmaceutical Industries and Associations, the Bill & Melinda Gates Foundation, the Global Health Drug Discovery Institute and the University of Dundee. The content of this publication only reflects the author’s views, and the JU is not responsible for any use that may be made of the information it contains.

RJP and CJJ received support from Australian Research Council Centre of Excellence for Innovations in Peptide and Protein Science.

This work made use of the Dutch national e-infrastructure with the support of the SURF Cooperative using grant no. EINF-2453, awarded to DLH.

SAKJ and VT acknowledge general financial support from the department of Chemical Biology and Drug Discovery at Utrecht University.

Utrecht Sequencing Facility is subsidized by the University Medical Center Utrecht, Hubrecht Institute, Utrecht University and The Netherlands X-omics Initiative (NWO project 184.034.019).

J.S. is funded by the Dutch Research Council NWO Gravitation 2013 BOO, Institute for Chemical Immunology (ICI; 024.002.009).

## Author contributions

Abbreviated author list:VT, DLH, OJDA, MAS, CF, AN, AA, NJM, DAAD, SWV, ML, WL, MC, TD, WD, ID, BJB, JS, ST, RJP, CJJ, FJMK, SAKJ

Conceptualization: FJMK, SAKJ

Methodology: VT, DLH, SAKJ

Validation: OJDA

Formal analysis: DLH, OJDA

Investigation: VT, DLH, OJDA, MAS, CF, AA, WL, NJM, MC, TD, WD, ID

Resources: VT, AN, DAAD, SWV, ML, WD, ID, BJB, RJP, CJJ, SAKJ

Data curation: DLH

Visualization: VT, DLH, MAS, SAKJ Funding acquisition: DLH, RJP, CJJ, FJMK

Project administration: DLH, FJMK, SAKJ

Supervision: DLH, BJB, JS, ST, RJP, CJJ, FJMK, SAKJ

Writing – original draft: VT, DLH, SAKJ

Writing – review & editing: VT, DLH, OJDA, MAS, CF, AN, AA, WL, NJM, DAAD, SWV, ML, MC, TD, WD, ID, BJB, JS, ST, RJP, CJJ, FJMK, SAKJ

## Competing interests

VT, DLH, FJMK and SAKJ are named inventors on a patent application that has been filed on 15 July 2022 entitled: Antiviral cyclic compounds (EP22185235; patent applicants: Universiteit Utrecht Holdings B.V. on behalf of Utrecht University). ID is an employee of Thermo Fisher Scientific. The remaining authors declare that they have no competing interests.

## Data and materials availability

High Throughput Sequencing data for the selections have been deposited in the DataverseNL repository at DOI https://doi.org/10.34894/WJRHLK.

The cryo-EM map of the SARS-CoV-2 spike ectodomain in complex with S1B3inL1 has been deposited to the Electron Microscopy Data Bank under the accession code EMD-16144. The corresponding atomic model has been deposited to the Protein Data Bank under the accession code 8BON. Materials generated in this study are available on reasonable request.

## Supplementary Materials

Materials and Methods

Figs. S1 to S20

Tables S1 to S5

References (*1–44*)

## Notes

### Summary of Updates

Minor wording changes in main tex; adding missing author.

