## Supporting information for "A broad-spectrum macrocyclic peptide inhibitor of the SARS-CoV-2 spike protein"

**This PDF file includes:**

### Materials and Methods

*Expression and purification of SARS-CoV-2 spike ectodomains*

To express the 2P-stabilized prefusion spike ectodomain for peptide selection, gene encoding residues 1 to 1200 of SARS2-S (GenBank: QHD43416.1) with proline substitutions at residues 986 and 987, a "AAARS" substitution at the furin cleavage site (residues 682 to 685), and residues 1 to 1160 of SARS-S (GenBank: AAP13567.1) with proline substitutions at residues 956 and 957, a C-terminal T4 fibritin trimerization motif, and a StrepTag were synthesized and cloned into the mammalian expression vector pCAGGS. Similarly, pCAGGS expression vectors encoding S1 or its subdomain S1B of SARS2 (S1, residues 1 to 682; S1B, residues 333 to 527) C-terminally tagged with the Fc domain of human or mouse IgG or Strep-tag were generated as described before.(*1*) Recombinant proteins were expressed transiently in FreeStyle 293-F Cells (Thermo Fisher Scientific) and affinity purified from the culture supernatant by protein A Sepharose beads (GE Healthcare) or Strep-Tactin beads (IBA) purification. For cryo-EM analysis, human codon-optimized gene was synthesized at Genscript encoding the 6P-stabilized SARS-CoV-2 S ectodomain expression construct(*2*) (S protein residues 1–1,213, Wuhan-Hu-1 strain: GenBank: QHD43416.1) with a C-terminal T4 foldon trimerization motif followed by an octa-histidine tag and a Twin-Strep-tag®.(*3*) The protein was expressed transiently in HEK-293T (ATCC® CRL-11268™) cells from pCAGGS expression plasmids, and secreted proteins were purified from culture supernatants using streptactin beads (IBA) following the manufacturer's protocol. Purity and integrity of all purified recombinant proteins were checked by Coomassie-stained SDS–polyacrylamide gel electrophoresis.

*Selection procedure*

Cyanomethyl esters of chloroacetylated D- and L-tyrosine (separate reactions) were charged with $tRNA_{CAU}$ as previously described.(*4*) Based on a previously reported method, selections were carried out against strep-tag immobilized protein target using MagStrep "type 3" XT Strep-Tactin coated magnetic beads (IBA lifesciences, Germany) for both D- and L-tyrosine initiated libraries in parallel. The DNA library, which encodes for 15 randomized NNK codons followed by a section encoding a CGSGSGS linker, was assembled by PCR, followed by transcription to RNA using T7 RNA polymerase (NEB) at 1mL overnight at 37 °C with 25 pmol DNA input and then purified by denaturing agarose gel electrophoresis. Puromycin ligation by T4 RNA ligase (NEB, USA), peptide translation using PURExpress (NEB, USA) and reverse transcription by ProtoScript II Reverse Transcriptase (NEB, USA) were all performed as reported previously.(*5*) Peptide libraries tagged with their encoding cDNA/mRNA were diluted with stock solutions of DPBS and BSA to achieve a final concentration of 1 X and 0.1%, respectively, with a total volume of 25 µl in the first selection round, and 14 µl in subsequent rounds. 1 µl of the reverse transcription reaction was diluted in 500 µl milliQ to serve as an input sample for qPCR analysis. The translation mixes were then sequentially incubated for three or six times with 2.5 µl of magnetic beads without target protein to pre-clear any peptides binding to the Strep-Tactin (omitted for the first round). The supernatants were then incubated with beads carrying protein target followed by stringent washing with 3 X 20 µl DPBS-T. Time and temperature of the incubation conditions are listed in Table S1. For the last two rounds of the selection against full-length SARS-CoV-2 spike, incubation with bovine serum was performed for 2.5 hours at 37 °C prior to incubation with beads. Bound peptides were then eluted by incubation with 50 µl MilliQ water for 5 minutes at 95 °C. The same wash and elution steps were performed with the third portion of pre-clear beads which was then used as a negative control sample in the qPCR analysis. Samples of the input, positive and negative (1 µl each in a 20 µL reaction) were analyzed by qPCR alongside a standard curve which was obtained by reverse transcription of the input library. Recoveries were calculated for the positive and negative samples by dividing the detected amount of DNA by the amount from the input sample, after compensating for dilution factors. The remaining solutions from the positive selections were amplified by PCR to act as input for a next round of selection, starting again with transcription to mRNA (without gel purification). This process was repeated until positive samples indicated enrichment above background. After enrichment was apparent, the output of round 3 or 5 was also used for sub-selection with additional targets. The DNA outputs of all rounds after the first were used for sequencing on the Illumina MiSeq platform using a 2 X 150 bp V2 reagent kit at the Utrecht UMC sequencing facility (USEQ).

*Solid Phase Peptide Synthesis*

Standard procedures are detailed below. After HPLC purification, all peptides used for cellular experiments were dissolved in 50 mM aqueous HCl and lyophilized for a total of three times to exchange the TFA salt of the peptide for HCl.

Standard peptide synthesis procedure (Microwave assisted SPPS)

All peptides shown in Table S2 were synthesized using micro-wave assisted Fmoc solid phase peptide synthesis on Rapp Polymere (Germany) tentagel S RAM resin on a Liberty Blue (CEM) Peptide Synthesizer at 25 µmol scale. Each coupling step of 4 minutes was performed with 5 equivalents of amino acid, 5 equivalents of Oxyma Pure and 10 equivalents of N,N'-diisopropylcarbodiimide (DIC) in DMF at 90 °C, with the exception of histidine couplings being performed at 50 °C for 10 minutes. Fmoc deprotection was performed using 20% piperidine in DMF for 1 min at 90 °C. Automated synthesis was followed by *N*-terminal chloroacetylation by using 5 equivalents of chloroacetic acid instead of amino acid in the relevant coupling step, or N-acetylation by treatment with 20% acetic anhydride in DMF for 2 minutes at 65 °C. Peptide cleavage, global deprotection, cyclization and subsequent purification were performed as reported before.(*6*) Peptide purity was analyzed by UV-HPLC using a 40 minute gradient from 100% buffer A (95% water, 5% Acetonitrile + 0.1% FA) to 70% buffer B (5% water, 95% Acetonitrile + 0.1% FA) with a Silicycle C18 column (Dr Maisch, Germany) (150 X 4.6 mm, 5 µm) and peptide identity verified by ESI-MS.

Dimeric peptide synthesis

Dimeric peptides were synthesized by standard automated Fmoc solid-phase synthesis on Rapp Polymere (Germany) tentagel XV-RAM resin using a Chorus (Gyros Protein Technologies, USA) system at 25 µmol scale. Branching was achieved using a $N^{\alpha,\varepsilon}$-bis-Fmoc-protected lysine amino acid, followed by coupling with 2-[2-[2-(Fmoc-amino)ethoxy]ethoxy]acetic acid to build up the PEG-like linker before synthesis of the peptide. A schematic representation of the synthesis is shown in Figure S19. Amino acid coupling reactions were performed using five equivalents of amino acid, five equivalents of Oxyma Pure, and ten equivalents of N,N'-diisopropylcarbodiimide (DIC) in DMF for 15 min at 55 °C in each cycle.This was followed by capping with 0.5 M each acetic anhydride and pyridine in DMF for 5 min at room temperature. Fmoc deprotection was achieved using 20% piperidine in DMF for 1.5 min at 80 °C. Peptides were then cleaved, cyclized and purified as above.1

*Protein Thermal Shift*

Purified peptides were prepared as a 400 µM stock solution in DMSO, as estimated by UV absorbance at 280 nm with calculated extinction coefficients (Expasy ProtParam, Swiss Bioinformatics Resource Portal). Thermal shift assay reaction mixtures for measurements with full length SARS-CoV-2 Spike contained 10 µM peptide, 0.25 mg/ml protein and 5x SYPRO orange dye (Thermo Fischer Scientific) in Dulbecco's phosphate-buffered saline (Gibco). Data collected for the S1b3inL1 peptide had an altered concentration of 0.1 mg/ml full length SARS-CoV-2 spike. Assays performed on SARS-CoV-2 RBD contained 2X SYPRO orange dye, 0.05 mg/ml protein for the S1b3inL1 peptide measurement and 0.1 mg/ml protein for the SARS2L1 peptide. An equal volume of DMSO compared to the peptide solution was added for the control experiments, leading to a final concentration of 5% DMSO. Thermal shift assays were performed on a Bio-Rad CFX96 PCR machine using the 'melting assay' protocol (FRET filter settings) with increment steps of 0.5 °C for 10 seconds from 20 to 75 °C. All measurements were performed in duplicate and averaged values are used to plot the final graphs shown. Temperature corresponding to the highest value of the d(RFU)/dT peak was used to determine the melting temperature ($T_m$).

*Pseudo-virus neutralization assay*

Human codon-optimized genes encoding the S proteins of SARS-CoV-2 S proteins corresponding to ancestral Wuhan-Hu-1 virus (GenBank: NC_045512.2) or VOCs Alpha (B.1.1.7), Beta (B.1.351), Gamma (P.1), Delta (B.1.617.2), and Omicron (B.1.1.529) were synthesized by GenScript. The production of SARS-CoV-2 S pseudo-typed VSV and the neutralization assay were performed as described previously.(*7*) Briefly, HEK-293T cells at 70 to 80% confluency were transfected with the pCAGGS expression vectors encoding SARS-CoV-2 S with a C-terminal cytoplasmic tail 18-residue truncation to increase cell surface expression levels. Cells were infected with VSV G–pseudo-typed VSVΔG bearing the firefly (Photinus pyralis) luciferase reporter gene at 48 hours after transfection. Twenty-four hours later, the supernatant was harvested and filtered through a 0.45-µm membrane. Pseudo-typed VSV was titrated on VeroE6 cells. In the virus neutralization assay, threefold serially diluted peptides were preincubated with an equal volume of virus at RT for 1 hour, then inoculated on VeroE6 cells, and further incubated at 37°C. After 20 hours, cells were washed once with PBS and lysed with Passive lysis buffer (Promega). The expression of firefly luciferase was measured on a Berthold Centro LB 960 plate luminometer using D-luciferin as a substrate (Promega). The percentage of neutralization was calculated as the ratio of the reduction in luciferase readout in the presence of peptide normalized to luciferase readout in the absence of peptide. The IC50 values were determined using four-parameter logistic regression (GraphPad Prism v8.3.0).

*Live virus neutralization assay*

The antiviral activity of Spike binding peptides against SARS-CoV2 was assessed in a live virus neutralization assay using the R-20 platform.(*8*) The compounds were initially diluted in cell culture medium (DMEM-5% FCS) to make 4x working stock solutions and then serially diluted further in the above media to achieve a 2-fold dilution series. On the day of the assay, HAT24 cells (Tea et al, PMID: 34228725) were trypsinized, stained with Nucblue in suspension (5% v/v) and then seeded at 16,000 cells in a volume of 40ul of DMEM-5% FCS per well in a 384-well plate (Corning #CLS3985). 20 uL of diluted compounds was added to the cells and the plates containing cells and compounds incubated for 1 hour at 37 °C, 5% CO2. 20 uL of virus solution at 8x103 TCID50/mL was then added to the wells and plates were incubated for a further 24 hours at 37 °C, 5% CO2. Stained cells were then imaged using the InCell 2500 (Cytiva) high throughput microscope, with a 10× 0.45 NA CFI Plan Apo Lambda air objective. Acquired nuclei were counted using InCarta high-content image analysis software (Cytiva) to give a quantitative measure of CPE. Virus inhibition/neutralization was calculated as %N= (D-(1-Q))x100/D, where; "Q" is the value of nuclei in test well divided by the average number of nuclei in untreated uninfected controls, and "D"=1-Q for the wells infected with virus but untreated with inhibitors. Thus, the average nuclear counts for the infected and uninfected cell controls get defined as 0% and 100% neutralization respectively. To account for cell death due to drug toxicity, cells treated with a given compound alone and without virus were included in each assay. The % neutralization for each compound concentration in infected wells was normalized to % neutralization in wells with equivalent amount of compound but without the virus to yield the final neutralization values for each condition. Dose response curves and interpolated IC50 values were determined using Sigmoidal, 4PL model of regression analysis in GraphPad Prism software (version 9.1.2, GraphPad software, USA).

*ELISA-based receptor binding inhibition assay*

The ACE2 receptor binding inhibition assay was performed as described previously.(*7*, *9*) Recombinant soluble ACE2 was coated on NUNC Maxisorp plates (Thermo Fisher Scientific) at 1 μg per well at RT for 3 hours. Plates were washed three times with PBS containing 0.05% Tween 20 and blocked with 3% BSA (Fitzgerald) in PBS containing 0.1% Tween 20 at 4°C overnight. Recombinant SARS-CoV-2 S RBD domain (200 nM) and serially diluted peptides or mAb REGN 10933 were mixed and incubated for 2 hours at RT. The mixture was added to the plate for 2 hours at 4°C, after which the plates were washed three times. Binding of SARS-CoV-2 S RBD domain to ACE2 was detected using 1:2000 diluted HRP-conjugated anti-StrepMAb (IBA) that recognizes the Strep-tag affinity tag on the SARS-CoV-2 S RBD domain. Detection of HRP activity was performed as described above (ELISA section).

*Cytotoxicity*

To indicate cell viability, an MTS, colourmetric assay was performed according to the manufacturer's protocol (Promega™ CellTiter 96® AQueous One Solution Cell Proliferation Assay). Briefly, VeroE6 cells were seeded in a 96 well plate format and peptide added to confluent cells with a total well volume of 100 µl in a concentration dependent manner, incubated at 37°C, for 20 hrs. Following incubation, the assay was completed by adding 20 µl of CellTiter 96® AQueous One Solution Reagent into each well and incubating at 37°C, for 2 hrs before recording the absorbance at 490nm using a 96-well ELISA plate reader (EL-808, BioTek). Cell viability was calculated as a % of the absorbance of the media only control, assuming cells were 100% viable.

*Hydrogen Deuterium Exchange Mass Spectrometry*

The SARS-S1B-2ST peptides were identified with 180 pmol diluted in a volume of 40 µl 100 mM Tris-HCl, 150 mM NaCl pH 7.5. To this mixture 20 µl of ice-cold 6 M Urea and 300 mM TCEP (pH 2.5) was added. The peptide identification measurements were performed in triplicate.

Two deuterated states were acquired; unbound state consisted of SARS-S1B and the binding state consisted of SARS-S1B and S1b3inL1. Binding took place with 3-fold excess of S1b3inL1 for one hour ice. Deuterium exchange reaction (Deuterium oxide 99.9% D, Sigma-Aldrich) proceeded on ice for 10 seconds, 1 min, 60 min and 240 min in 95% D2O with final concentration of 100 mM Tris-HCl, 150 mM NaCl and 1.0 mM EDTA pH 7.5. Reaction was quenched with ice-cold 6 M urea, 300 mM TCEP at a final pH of 2.5. Each time point was acquired in quadruplicate.

The samples were immediately injected into a HDX/nano-Acquity UPLC system (Waters) coupled to a Xevo QTof G2 instrument (Waters). The system was equipped with a pepsin column (NovaBioassay, immobilized pepsin column) at 20°C, the peptides were trapped in-line for 3 min at 125 µl/min with 0.1% formic acid in MQ with pH set on 2.5 on a Acquity UPLC C18 vanguard pre-column 300 A 1.7 µM 2.1 mm at 0.5 °C. The peptides were separated on the Waters BEH C18 column (Xbridge BEH300 C18 3.5 µM 1.0x150 mm) using a 15 min gradient (8 - 85% B). Formic acid 0.1% (v/v) in MQ was used as mobile phase A and 0.1 % (v/v) in acetonitrile was used as mobile phase B.

Acquired data from the peptide identification measurements were processed with ProteinLynx Global Server (3.0.1) software. The PLGS files were analyzed in Dynamx 3.0 from Waters (minimum intensity 100, minimum sequence length 4 and file threshold 2). The acquired mass spectra were lock mass corrected with Leu-enkephalin (161 of 440 unique peptides survived filtering). Subsequently, in DynamX the deuterium uptake was calculated by difference in uptake between the unbound SARS-S1B state and binding state with

S1b3inL1. Statistical analysis was performed in Excel (Microsoft) with an independent t-test, peptides that showed a significant difference were filtered with a $p < 0.001$.

*Cryo-electron microscopy sample preparation and data collection*

To prepare the spike-peptide complex for cryo-EM analysis, 5.4 µl of SARS-CoV-2 hexaproline S-ectodomain, at a concentration of 28 µM (based on the molecular weight of the spike protomer) was combined with 0.6 µl of 1.1 mM S1B3inL1 and incubated for ~10 min at room temperature. Immediately before blotting and plunge freezing, 1 µl of 0.07% (w/v) fluorinated octyl maltoside (FOM) was added to the sample, resulting in a final FOM concentration of 0.01% (w/v). Subsequently, 3 µl of spike-Fab complex was applied to glow-discharged (20 mAmp, 30 sec, Quorum GloQube) Quantifoil R1.2/1.3 grids (Quantifoil Micro Tools GmbH), blotted for 5 s using blot force 0 and plunge frozen into liquid ethane using Vitrobot Mark IV (Thermo Fisher Scientific). The data were collected on a Thermo Scientific™ Krios™ G4 Cryo Transmission Electron Microscope (Cryo-TEM) equipped with Selectris X Imaging Filter (Thermo Fisher Scientific) and Falcon 4 Direct Electron Detector (Thermo Fisher Scientific) operated in Electron-Event representation (EER) mode. In total, 2612 movies were collected at a nominal magnification of 130,000×, corresponding to a calibrated pixel size of 0.929 Å/pix over a defocus range of −0.75 to −1.5 µm. A full list of data collection parameters can be found in Table S4.

*Single-particle image processing*

Collected movie stacks were imported into Relion version 3.1.3.(*10*) Drift and gain correction were performed using Relion's implementation of MotionCor,(*11*) and CTFFIND-4.1(*12*) was used to estimate the contrast transfer function for each movie. Movies with a CTF-estimated resolution of worse than 5 Å were discarded, leaving 2576 micrographs for further processing. Template-free auto-picking with a Laplacian-of-Gaussian filter was then used to select 484,282 particles. Fourier binned (4 × 4) particles were extracted in a 100-pixel box and subjected to a single round of 2D classification, ignoring CTFs until the first peak, after which 368,031 particles were retained. Using the 'molmap' command in UCSF chimera,(*13*) a SARS-CoV-2 spike structure (PDB ID: 6VSB)(*14*) was used to generate a 50 Å resolution starting model for 3D classification. Particles selected from 2D classification were subject to a single round of 3D classification (with C1 symmetry). Particles corresponding to a single, well-defined class (91,707 particles) with two open RBDs were re-extracted in a 300-pixel box. During extraction, particles were Fourier binned by a non-integer value, resulting in a final pixel size of 1.2384 Å. Subsequent 3D auto-refinement (with C1 symmetry) and post-processing yielded a map with a resolution of 3.72 Å. Relion's Bayesian polishing procedure was then performed on these particles, with all movie frames included, which improved the resolution to 3.18 Å. Subsequently, CTF refinement was performed to estimate anisotropic magnification, beamtilt, trefoil, $4^{th}$ order aberrations, per-particle defocus and per-micrograph astigmatism(*15*). Collectively, this improved the resolution to 3.09 Å, based on the gold-standard FSC = 0.143 criterion. To improve the quality of the S1B3inL1 density, a focused 3D classification approach was employed. A soft mask was placed over the map to isolate the peptide-bound, closed RBD, and the particles were subjected to masked 3D classification without alignment using a regularization parameter ('T' number) of 60, over

100 iterations. Classes were inspected visually, and 38,457 particles from the best resolved map (with assigned orientation information) were used to generate half maps using the relion_reconstruct tool. Post-processing yielded a map with a resolution of 3.2 Å. Local resolution estimations were performed using Relion. An overview of the data processing pipeline is shown in Supplementary Fig. 2.

Model building and refinement.

UCSF Chimera(*13*) (version 1.15.0) and Coot(*16*) (version 0.9.6) were used for model building. The structure of the SARS-CoV-2 spike glycoprotein in complex with the 87G7 antibody Fab fragment (PDB ID 7R40)(*17*) was used as a starting point for modelling the S1B3inL1-bound spike, and RBD crystal structure (residues 333-526; PDB ID 6M0J(*18*) (53)) were individually rigid-body fitted into the density map using the UCSF Chimera "Fit in map" tool. As a starting point for modelling the S1B3inL1 peptide, AlphaFold2(*19*, *20*) was used to generate five models of S1B3inL1. To promote cyclisation, an N-terminal cysteine was added to the peptide sequence. The predicted model which most closely resembled the experimental density was rigid body fitted into the EM density map using the UCSF Chimera 'fit in map' tool and then combined with the spike model. The resulting model was then edited in Coot using the 'real-space refinement, carbohydrate module(*21*) and 'sphere refinement' tool. Subsequently, ISOLDE(*22*) was used to perform molecular-dynamics flexible fitting on S1B3inL1 and the surrounding residues. Following this, iterative rounds of manual fitting in Coot and real space refinement in Phenix(*23*) were carried out to improve non-ideal rotamers, bond angles and Ramachandran outliers. During refinement with Phenix, secondary structure and non-crystallographic symmetry restraints were imposed. The final model was validated with MolProbity,(*24*) EMRinger(*25*) and Privateer (glycans).(*26*, *27*)

Structure analysis and visualization.

Spike residues interacting with S1B3inL1 were identified using PDBePISA(*28*) and LigPlot+.(*29*) Figures were generated using UCSF ChimeraX.(*30*) Structural biology applications used in this project were compiled and configured by SBGrid.(*31*)

*Dataset collection for sequence alignment*

All protein sequence records belonging to all lineages of the Coronaviridae family (approximately 4000000 protein sequences) were retrieved from the NCBI virus dataset. All sequences with annotations related to the spike protein were extracted and redundancy was removed to a similarity threshold of 99% sequence identity over the full spike protein sequence.(*32*) All ambiguous sequences (defined as containing at least a single unresolved character) and sequences shorter than 1000 amino acids in length were removed, yielding a non-redundant dataset of 725 spike protein sequences. This dataset was aligned using the default parameters of the MAFFT-ginsi.(*32*, *33*)

To analyze variability among redundant SARS-COV2 spike sequences, the initial redundant dataset from NCBI was aligned in UPP(*34*) once ambiguous sequences were removed. A guide tree for the ultra-large alignment was inferred from the non-redundant spike dataset (725 sequences) using the default parameters of IQ-TREE.(*34, 35*) The non-redundant spike dataset also served as a seed alignment for the redundant dataset.

*Phylogenetic reconstruction*

The non-redundant coronaviridae phylogeny was inferred by maximum likelihood (ML) in IQ-TREE.(*35*) The ML sequence evolution model (WAG+F+R10) was selected Bayesian and Akaike information criteria, as implemented in ModelFinder.(*36*) Branch supports were measured by ultrafast bootstrap approximation and approximate likelihood ratio test, each conducted to 1000 replicates in IQ-TREE.(*37*) 100 independent replicates of tree-search were conducted, 1 of which was rejected by the approximately unbiased test conducted to 10,000 replicates.(*38*) The topology that is presented in Figure S17 was chosen for subsequent analysis from the remaining 99 topologies as it scored the highest likelihood and had high branch supports at key bifurcations in coronaviridae evolution.

*Molecular dynamics simulation*

Atom coordinates not modeled into the CryoEM density were transplanted from a complete spike model (PDB: 6VSB) accessed from the CHARMM GUI covid-19 database.(*39*) Site-specific N-linked complex glycans (66 in total, 22 per monomer) were added as covalent adducts at experimentally identified sites(*40*) using the CHARMM GUI online portal, which was also used to generate the system topology files.(*39*) Parameters for the S1B2inL1 peptide were generated in CHARMM General Force Field (CGenFF), once explicit hydrogen atoms were added.(*41*)

All-atom molecular dynamics simulations were performed in the GROMACS MD engine(*42*) using the CHARMM36M force field.(*43*) MD integration used a time step of 2 fs. Columb interactions were modeled using the particle mesh Ewald (PME) algorithm with a radius of 1.2 nm and Leonard Jones interactions used a cut-off scheme and a radius of 1.2 nm. Hydrogen bonds were constrained using the LINCS algorithm.(*44*) 6 independent replicates of equilibration (from the NVT ensemble onwards) and production MD were performed. The spike trimer glycoprotein and S1b3inL1 peptide complex was put in a rhombic dodecahedral box extending 1 nm in all directions from the boundary to the closest atom in the box. The box was filled with 202843 TIP3P water molecules and the charge was neutralized with sodium and chloride ions. To replicate physiological conditions, the sodium chloride concentration of the system was set at 150 mM. The system was minimized by steepest descent with a step size of 0.01 nm and convergence criteria of $F_{max}$ of 1000 KJ/mol/nm. The system was equilibrated in the NVT ensemble for 100 ps to 300 K with a velocity rescale thermostat and harmonic restraints on all heavy atoms. To avoid instabilities, solvent and ions were coupled independently of the protein, peptide and glycans. The system was brought to 1 Bar pressure in the NPT ensemble using a Berendsen barostat for 100 ps with harmonic restraints on all heavy atoms. Following temperature and pressure equilibration, unrestrained production MD was performed for 300 ns in the NPT ensemble using a velocity rescale thermostat and a Parrinello-Rahman barostat with references of 300

K and 1 Bar, respectively. Trajectories were analyzed using the MDTraj python package and every 10th frame was sampled.

*Surface plasmon resonance*

SPR measurements were performed using a Biacore™ T200 (GE Healthcare) with 10 mM HEPES pH 7.5, 150 mM NaCl, 0.005 vol% Tween® 20 and 0.1 vol% DMSO as running buffer. The sensorchip surface of a CM5 chip (GE Healthcare) was activated using NHS/EDC before immobilisation of SARS-CoV-2 spike protein variants in 10 mM acetate buffer pH 4.0. All experiments were conducted at 25 °C in single cycle kinetics mode with a flow rate of 50 µL min-1. A concentration series of 1-500 nM of peptide was injected over captured protein to measure association and dissociation. Dissociation constants (KD) were derived from reference flow cell and blank-subtracted sensorgrams and analysed using the Biacore™ Insight Evaluation Software.

**Figs. S1 to S20**

*Protein immobilization*

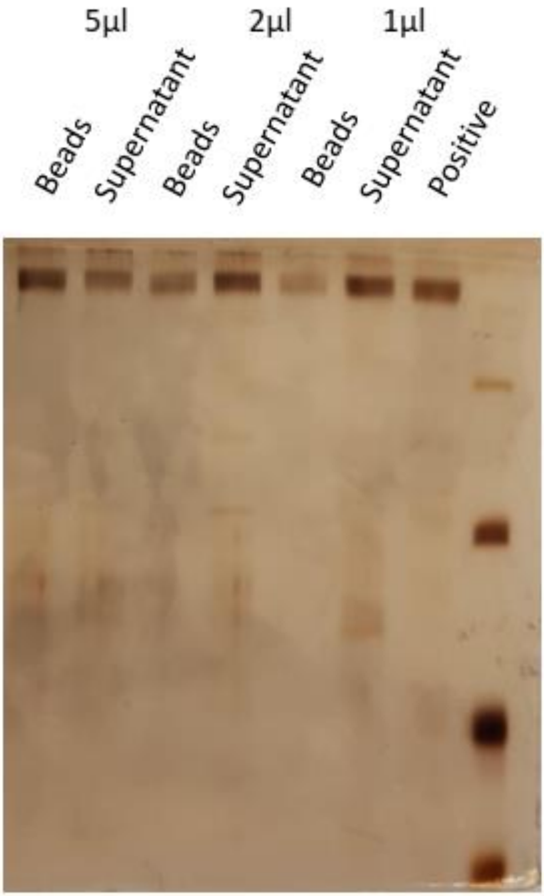

Fig S1. Optimization of bead to protein ratio for immobilization of SARS-CoV-2 spike protein (200 ng) on various volumes of magnetic Streptactin beads, as indicated. Protein visualized by silver staining.

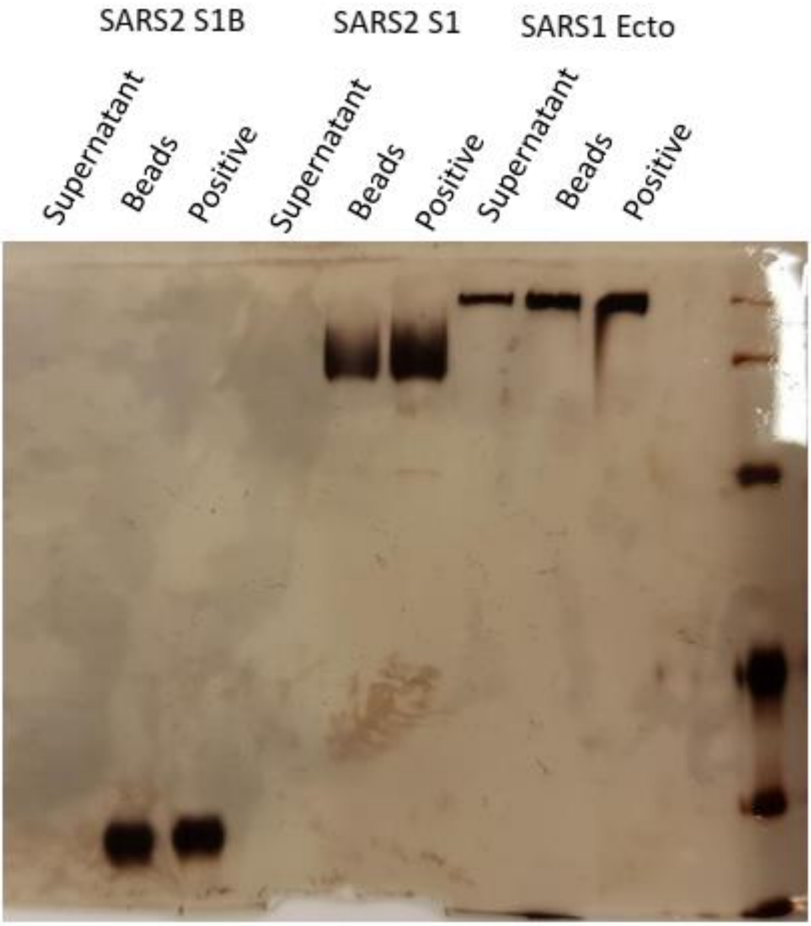

Fig S2. Verification of immobilization of other protein targets, with 200 ng indicated domain on 5 μL magnetic Streptactin beads. Protein visualized by silver staining.

*Selection*

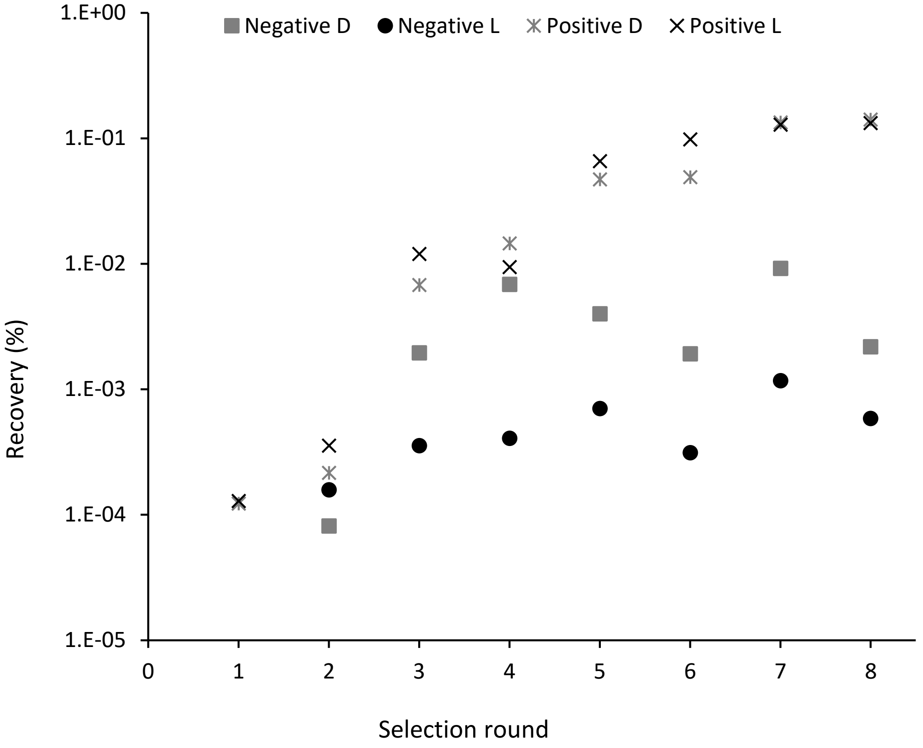

Figure S3 Log transform for DNA recovery across selections rounds, representing the percentage of binding peptides in the library against immobilized protein target on beads (positive) or beads alone (negative) for both L- and D-tyrosine initiated libraries. Rounds where the library was incubated with fetal calf serum are indicated with *.

2

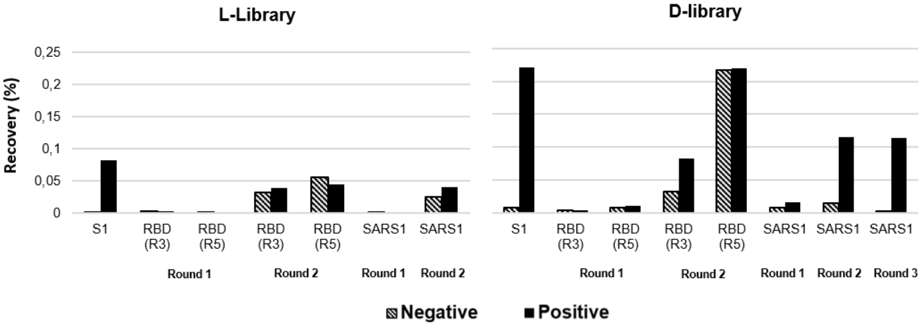

Figure S4 DNA recovery for sub-selections with both d- and l-tyrosine initiated libraries against different domains of SARS-CoV-2 spike protein or against SARS1-S ectodomain. Positive indicates recovery against immobilised target, whereas negative is the recovery against immobilisation beads alone. Input used was from round 5 of the initial selection against full-length SARS-CoV-2 spike, except for the S1B domain where input round is indicated in parentheses.

*Sequence analysis*

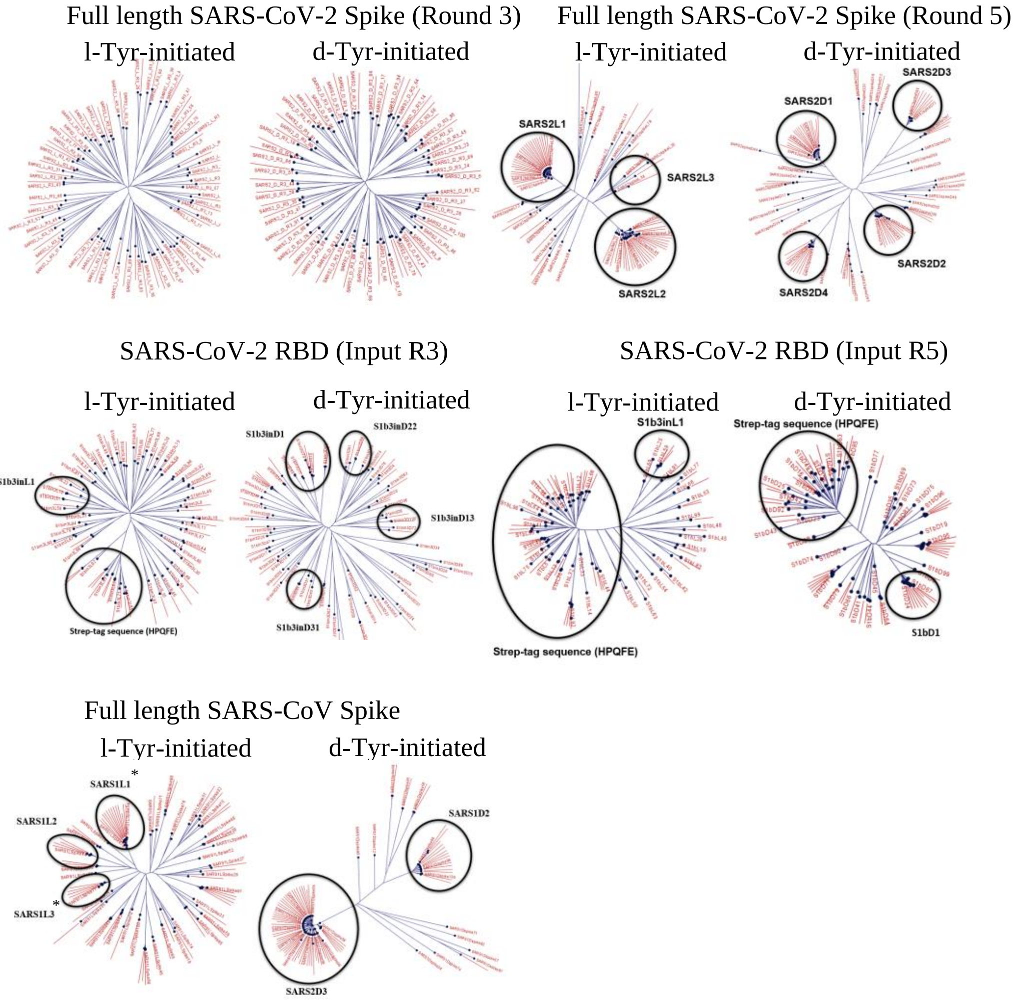

Fig. S5. Phylogenetic tree representation of the high-throughput sequencing results for different rounds of the selection, based on a neighbour-joining algorithm following multiple sequence alignment of unique sequences. Clusters that correspond to peptides chosen for further investigation are indicated with circles. * denotes sequences that were unsuccessful on SPPS.

*Peptide characterization*

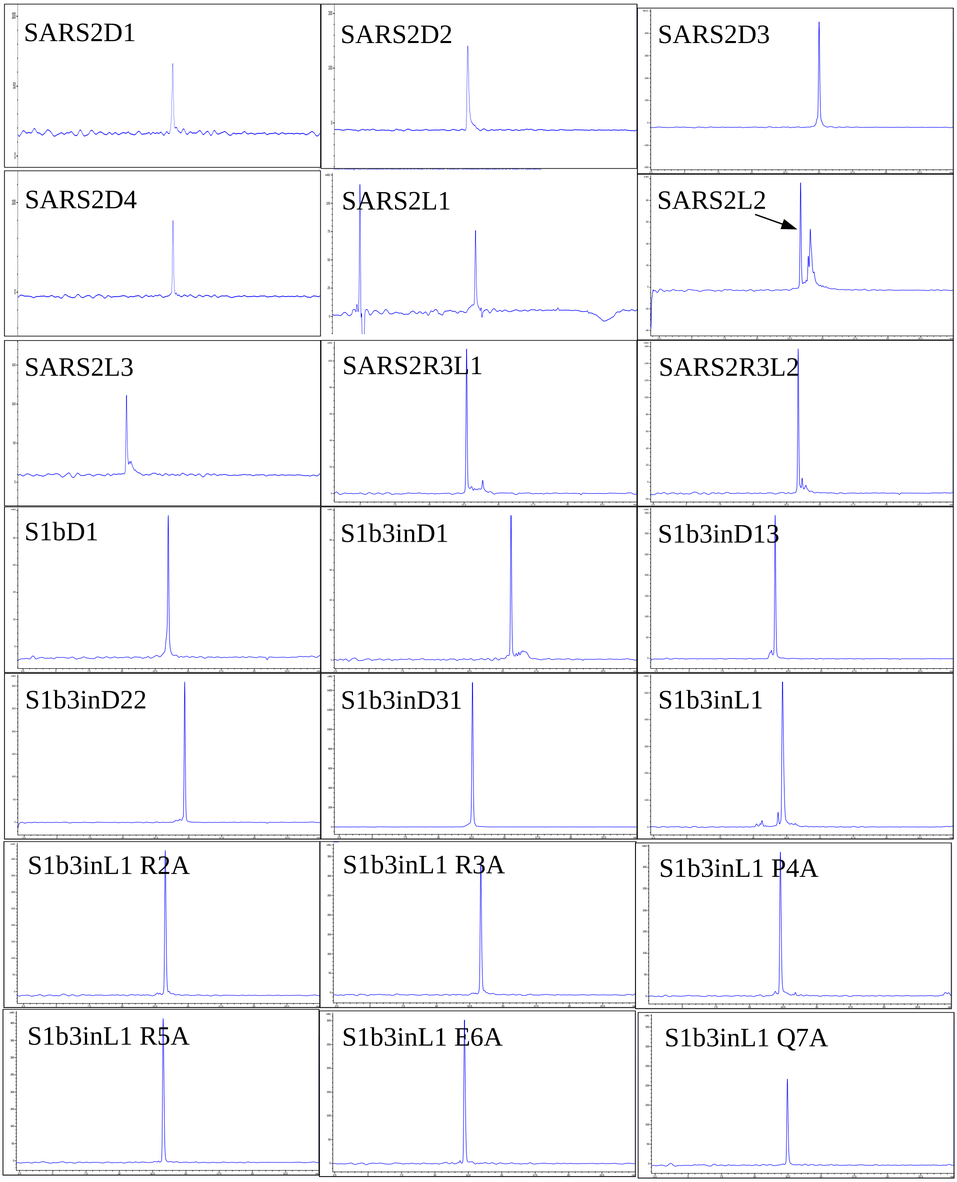

Figure S6. HPLC analysis of purified peptides. UV-absorption traces at 215 nm. The arrow indicates desired peak where substantial contaminants remain after the first purification run.

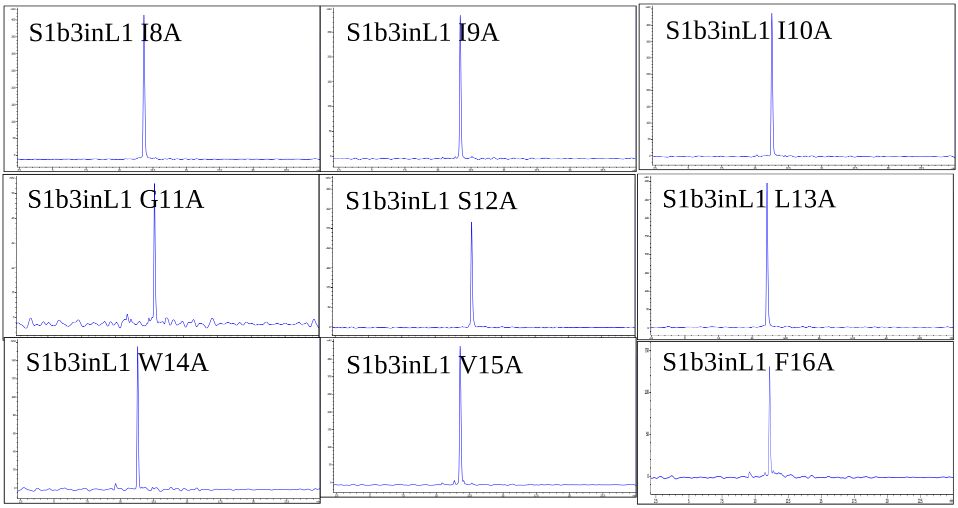

Figure S6 contd. HPLC analysis of purified peptides. UV-absorption traces at 215 nm.

*Activity testing*

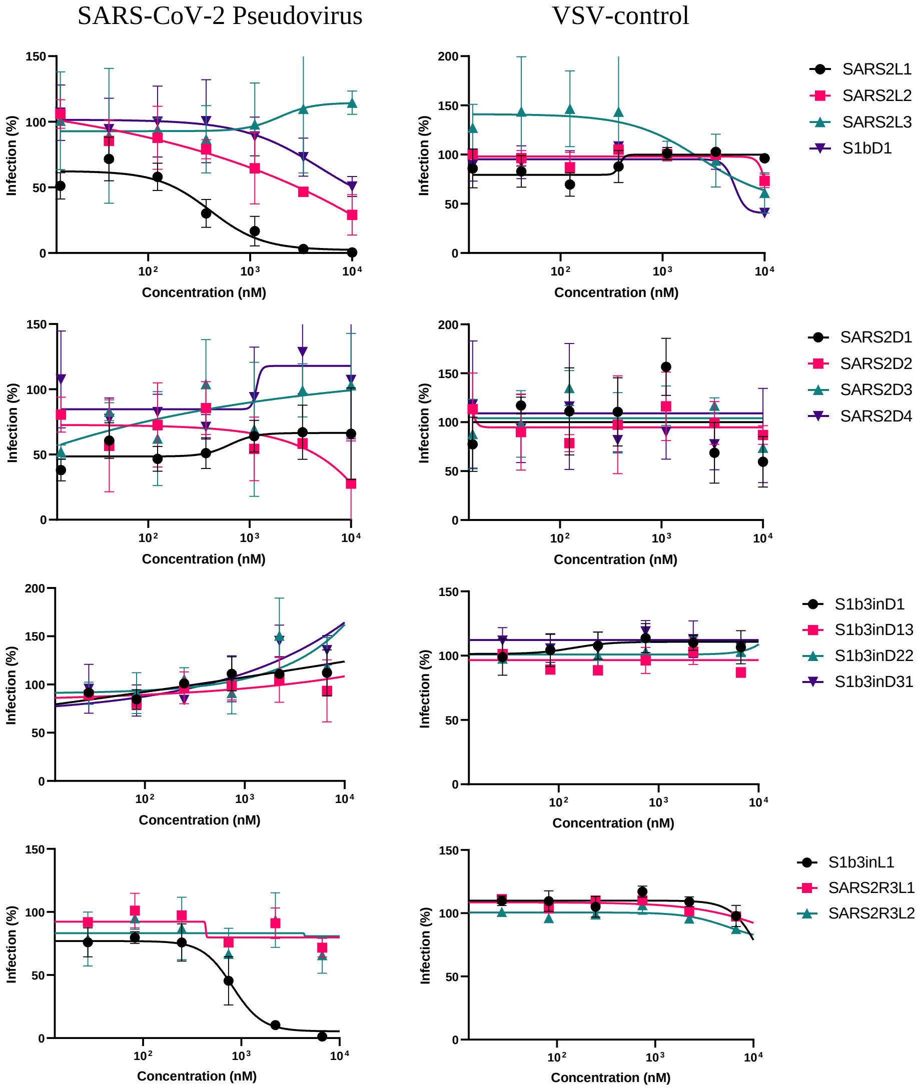

Figure S7. Luciferase-encoding pseudovirus neutralization assay of peptides against either SARS-CoV-2 VSV or VSV control. A three-fold dilution of peptide was performed starting at 10 µM. Peptides SARS2L1 and S1b3inL1 showed a clear concentration-dependent inhibition against the SARS-CoV-2 VSV but not against the control.

*Thermal shift assays*

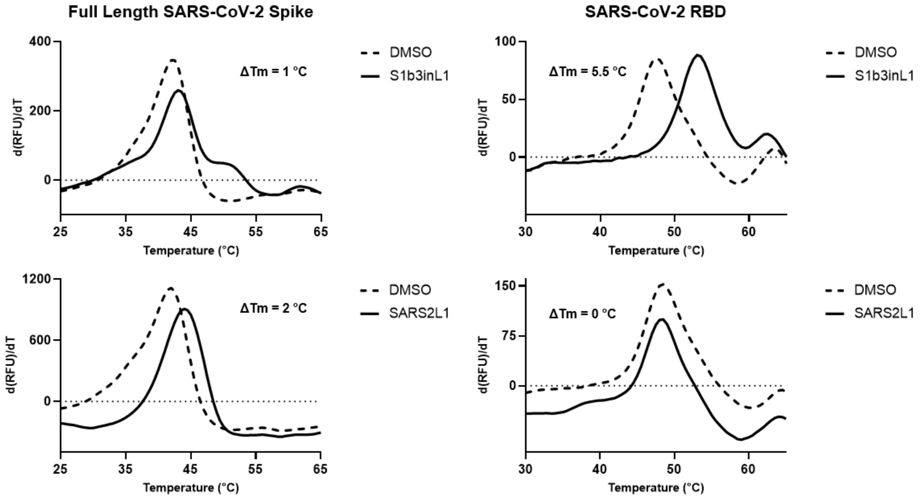

Figure S8. Thermal shift analysis of peptides S1b3inL1 and SARS2L1 against either full length SARS-CoV-2 or isolated RBD. Change in melting temperature (ΔTm) was observed for both peptides against full length protein, where only S1b3inL1 showed an increase against RBD.

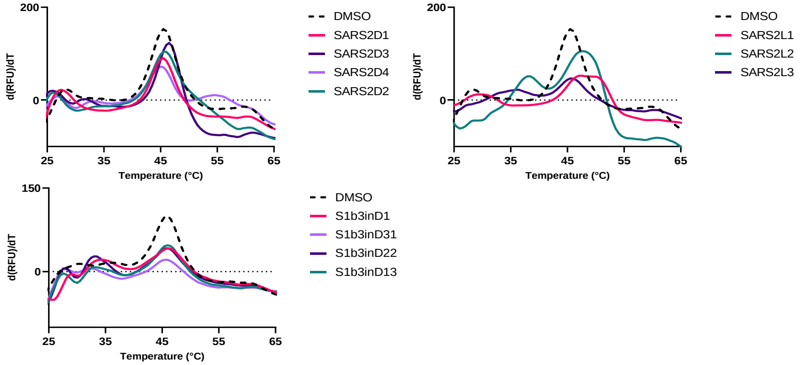

Figure S9. Thermal shift analysis of non-inhibitory peptides in fig. S7 tested against either full length SARS-CoV-2 or isolated RBD. Change in melting temperature (ΔTm) was observed for both peptides against full length protein, where only S1b3inL1 showed an increase against RBD.

*Toxicity*

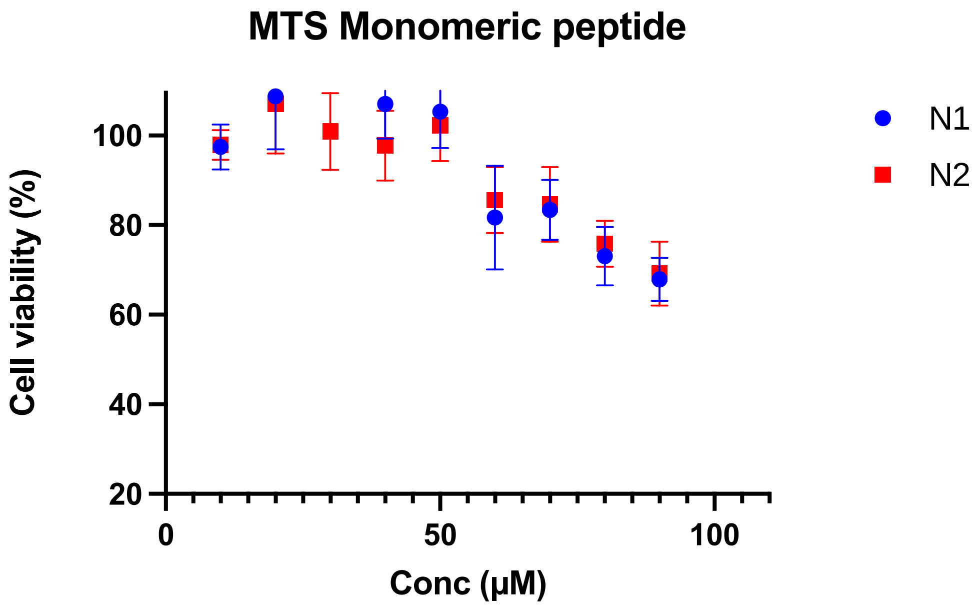

Fig S10. Toxicity assay for monomeric S1b3inL1 peptide

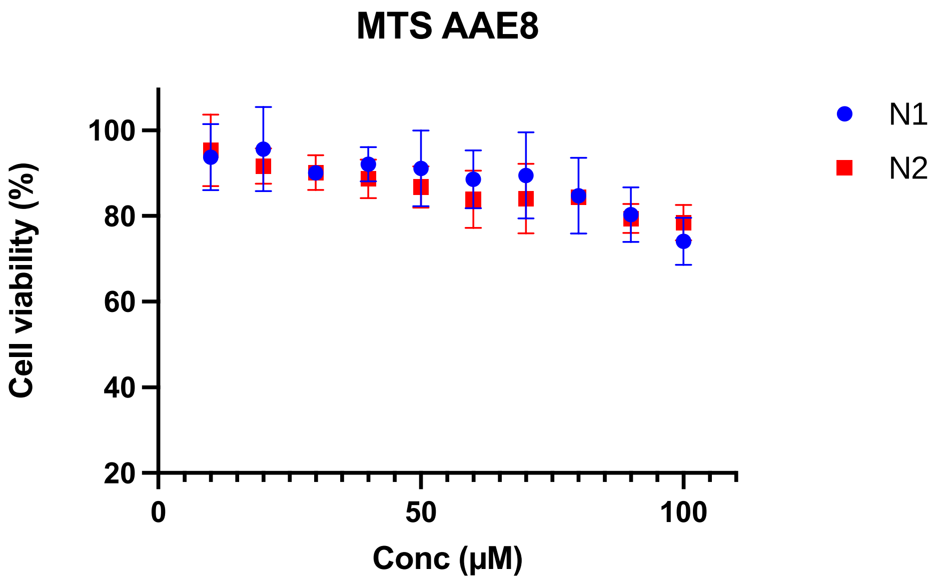

Fig S11. Toxicity data for S1b3inL1 AEEA8 dimer

*Hydrogen-deuterium exchange*

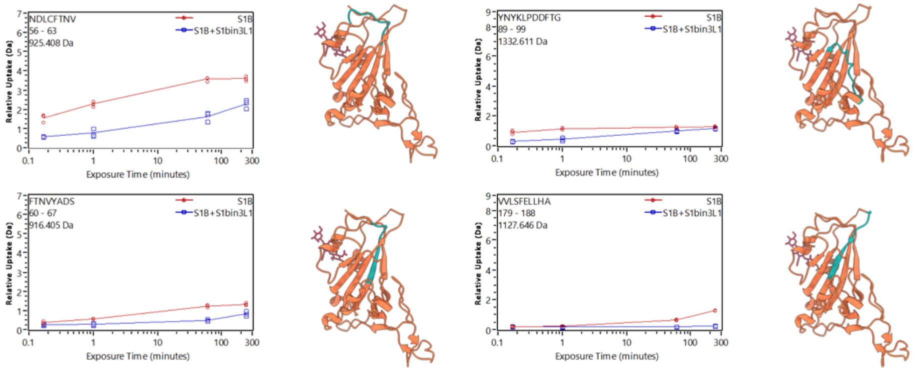

Figure S12. Binding epitope on SARS-CoV-2 RBD of S1b3inL1 elucidated by HDX. The absolute deuterium uptake in Dalton for different peptide fragments in the HDX experiment is shown. Individual data points (N=4) are plotted. Next to each graph, a cartoon representation of the SARS-CoV-2 RBD (coral) is shown with the identified peptide colored cyan.

*Cryo-EM3*

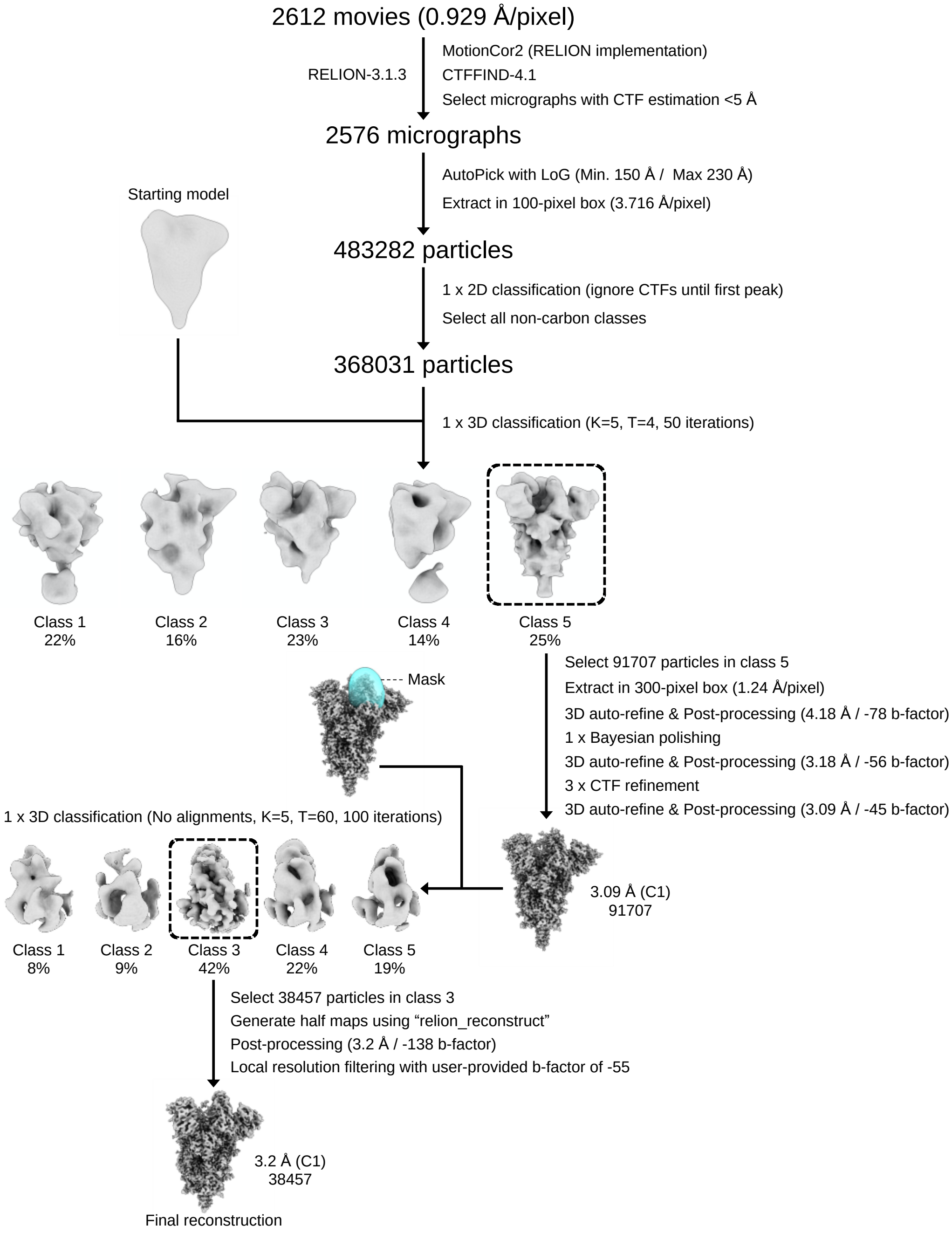

Figure S13. Cryo-EM data processing pipeline SARS-CoV-2 S ectodomains in complex with S1BinL1.

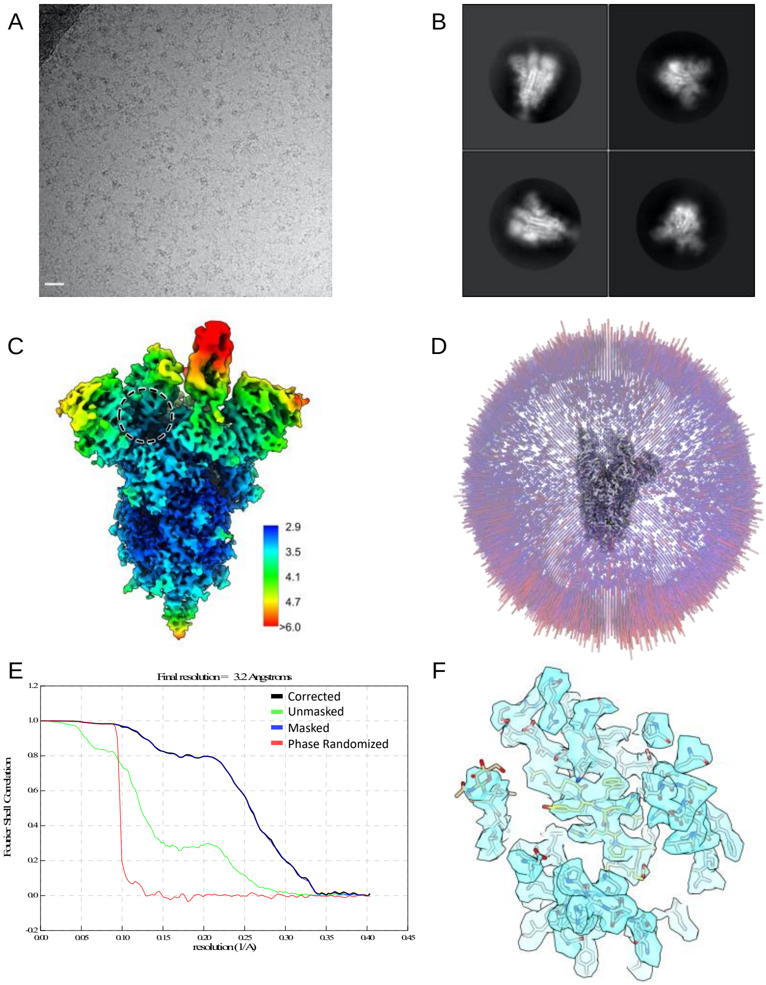

Figure S14. Cryo-EM data processing and validation of the SARS-CoV-2 spike ectodomain in complex with S1B3inL1. A. Representative electron micrograph of SARS-CoV-2 S ectodomains in complex with S1B3inL1, embedded in vitreous ice. Scale bar: 250 Å. B. Representative 2D class averages generated from the final particle stack. C. Local resolution filtered EM density map for the C1 refined SARS-CoV-2 spike in complex with S1B3inL1, colored according to local resolution which was calculated in Relion-3.1.3. The macrocyclic peptide binding site is circled. D. Angular distribution plot of the final C1 refined EM density map. E. Gold-standard Fourier shell correlation (FSC) curve generated from the independent half maps contributing to the 3.2 Å global resolution density map. F. EM density and fitted atomic coordinates for the S1B3inL1 binding site on the closed RBD. The spike protomers are colored blue, grey and pink, and the macrocyclic peptide and glycans are colored gold and tan, respectively.

*Molecular dynamics*

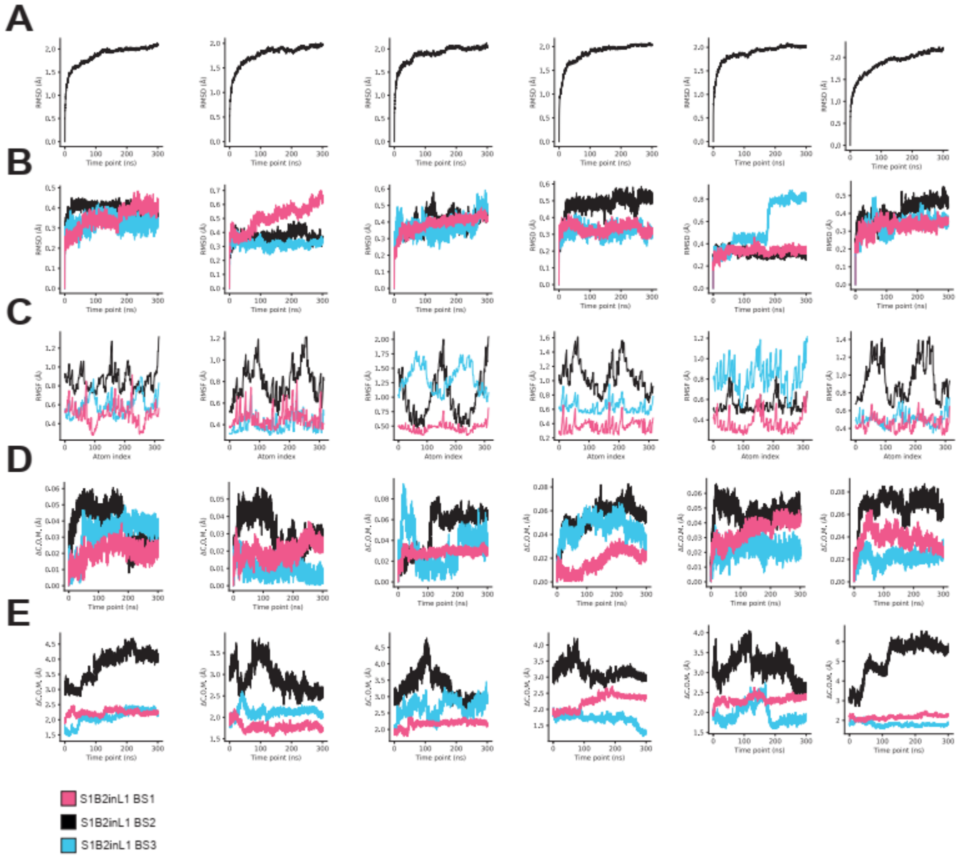

Figure S15. Replicate simulation data. **A**. RMSD of alpha carbons over 300 ns ordered in replicates 1-6 (left to right). Each system is slow to equilibrate, requiring 100 - 250 ns of sampling. **B**. RMSD over all atoms for S1b3inL1 in each of three binding sites (BS1 - BS3). **C**. RMSF for over all atoms in S1b3inL1 in each of three binding sites. BS1 (magenta) is more conformationally stable than the alternate binding poses (blue, black). **D**. Centre of mass (C.O.M.) deviation for each of the 3 peptides from the initial binding pose. **E**. C.O.M difference between each peptide and its closest RBD in the initial pose. BS1 exhibits motions coupled to the S1B domain, whereas BS2 demonstrates C.O.M. movements independently of the S1B domain.

*Sequence conservation*

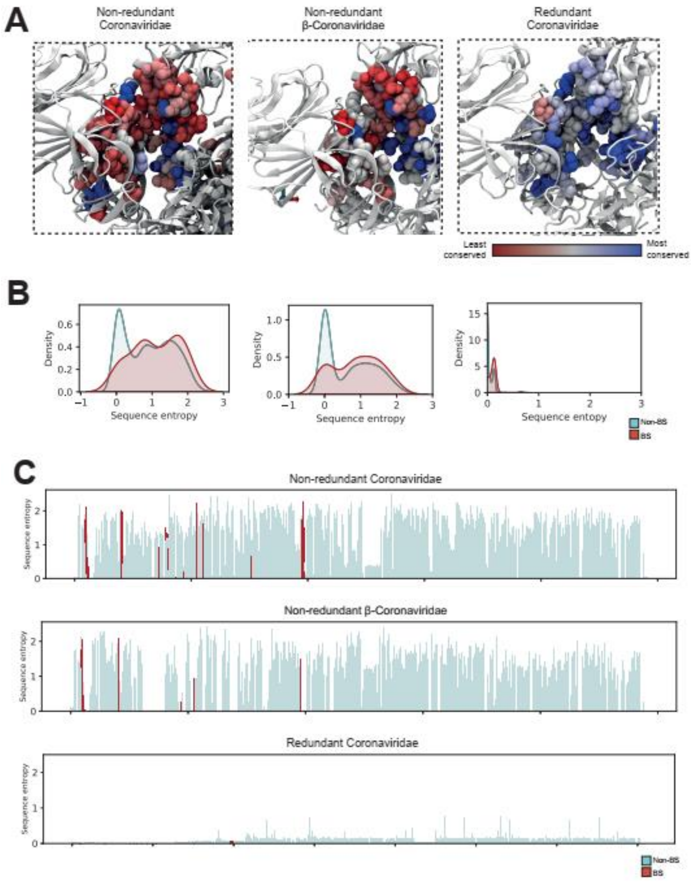

Figure S16. Sequence entropy analysis of the peptide-binding site across coronaviruses. **A.** Conservation of the peptide-binding site across all non-redundant coronavirus spike protein sequences (left); all non-redundant β-coronavirus spike protein sequences (middle); all redundant coronavirus spike sequences (right). Residues that interact with the peptide are shown as VDW spheres and are coloured by relative sequence entropy. **B**. Kernel density estimate plots of binding site residue entropy (maroon) and non-binding site residue entropy (cyan) calculated over all non-redundant coronavirus spike sequences (left); all non-redundant β-coronavirus spike protein sequences (middle); all redundant coronavirus spike sequences (right). **C**. Site-wise sequence entropies for the full-length spike protein calculated over all non-redundant coronaviridae spike sequences (top); all non-redundant β-coronavirus spike protein sequences (middle); all redundant coronavirus spike sequences (bottom). The peptide-binding site is similarly conserved to most other spike protein residues, excluding those that are stringently conserved to maintain the spike structure and function when investigated over multiple tiers of phylogenetic inclusion (all coronavirus spike proteins, β-coronavirus spike proteins, and only SARS-COV2 spike proteins).

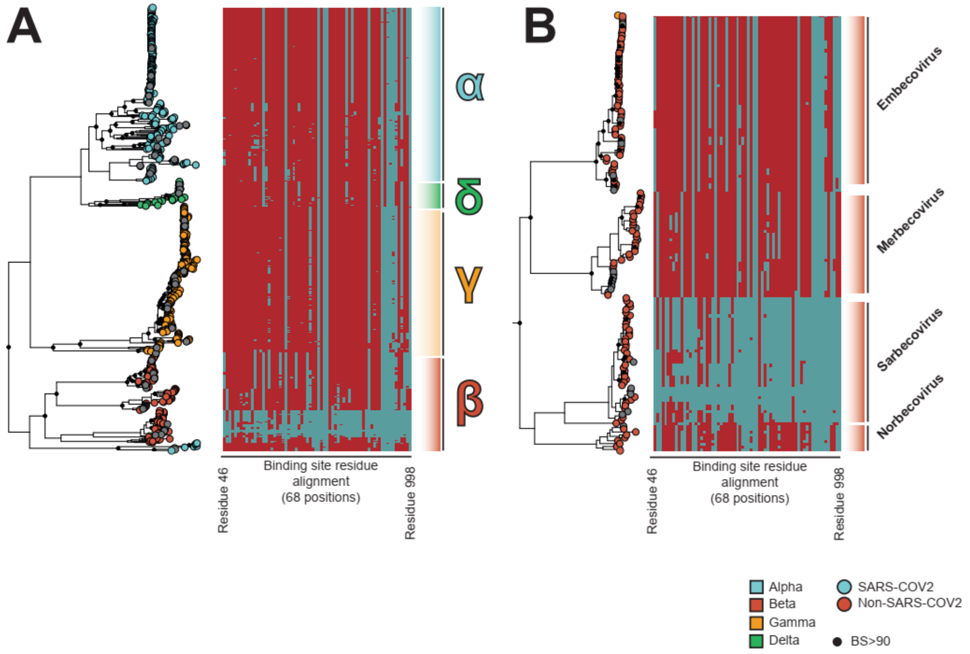

Figure S17. Phylogenetic analysis of coronaviridae spike proteins. **A**. Maximum likelihood phylogenetic reconstruction of all coronaviridae spike proteins. Major lineages (α, β, γ and δ) are highlighted and labeled. The topology is rooted at the midpoint and is reconstructed from 725 non-redundant amino acid sequences. Peptide binding site residues (designated as SARS-COV2 residue, cyan, or non-SARS-COV2 residue, maroon) are aligned respective tips. **B**. Clade view of the β-coronavirus lineage presented in panel A. Major β-coronavirus lineages are labeled. The peptide-binding site is marginally conserved across non-SARS-COV2 sarbecoviruses and norebecoviruses, but is lost in merbecovirus, embecovirus and α-, γ-, δ-coronavirus lineages.

*Surface plasmon resonance*

a)

| | $K_d$ (nM) | | | | $k_{on}$ ($M^{-1}.S^{-1}$) | | | | $k_{off}$ ($S^{-1}$) | | | |
|---|---|---|---|---|---|---|---|---|---|---|---|---|
| | WT | Alpha | Beta | Delta | WT | Alpha | Beta | Delta | WT | Alpha | Beta | Delta |
| **S1b3inL1** | 46 | 39 | 60 | 50 | 3.12 E+5 | 8.54 E+5 | 1.49 E+5 | 1.62 E+5 | 1.44 E-2 | 2.91 E-2 | 9.61 E-3 | 7.29 E-3 |
| **S1b3inL4** | 206 | 95 | 135 | 115 | 1.34 E+5 | 2.82 E+5 | 1.50 E+5 | 2.40 E+5 | 2.77 E-2 | 2.68 E-2 | 2.03 E-2 | 2.76 E-2 |
| **SARS2L1** | 0.8 | 0.65 | 1.1 | nd | 1.20 E+6 | 1.42 E+6 | 1.01 E+6 | nd | 1.20 E-3 | 1.17 E-3 | 1.22 E-3 | nd |

b)

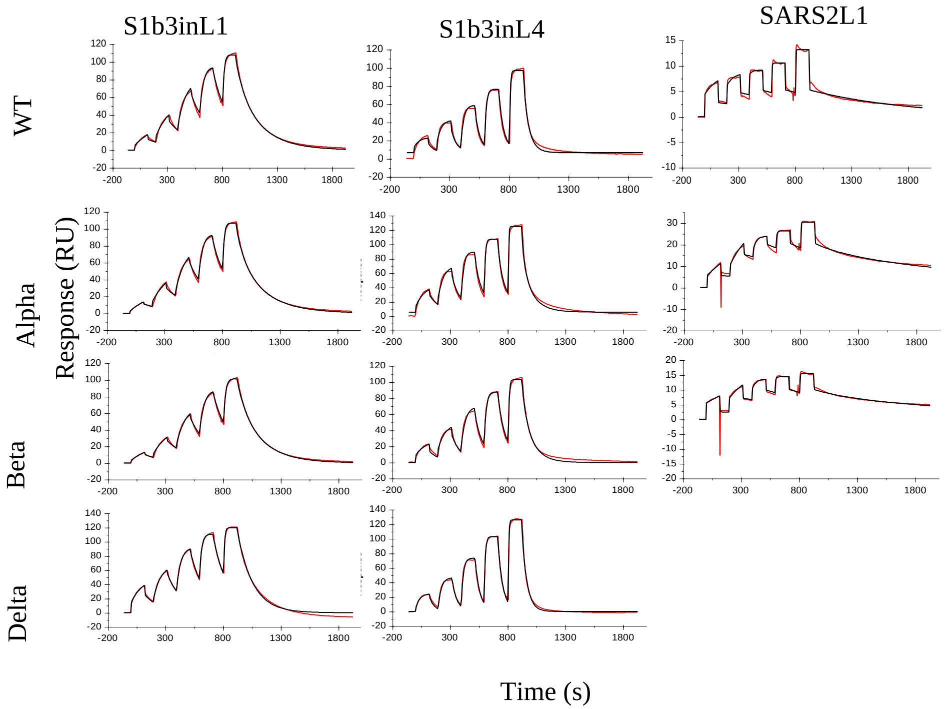

Figure S18 (a) Binding kinetics and dissociation constants ($K_d$) as determined by SPR for peptides (S1b3inL1, S1b3inL4 and SARS2L1) against either WT SARS-CoV-2 spike or mutants. Not determined values are denoted as nd. (b) Representative SPR sensorgrams showing the binding of different peptides to the different variants of SARS-CoV-2 spike protein. Fit (black) of the data (red) is shown using a 1:1 binding model.

*Dimer peptides*

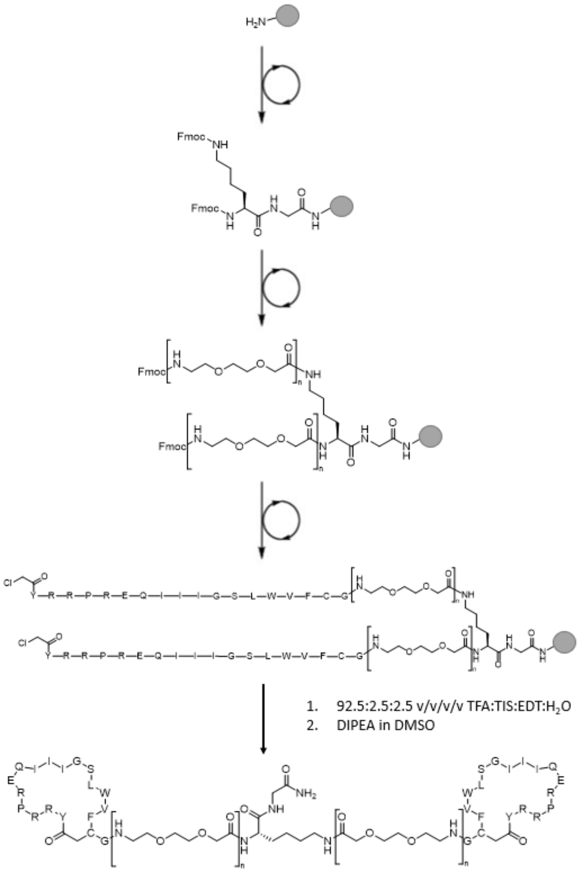

Figure S19 General synthesis strategy for S1b3inL1 dimer peptides.

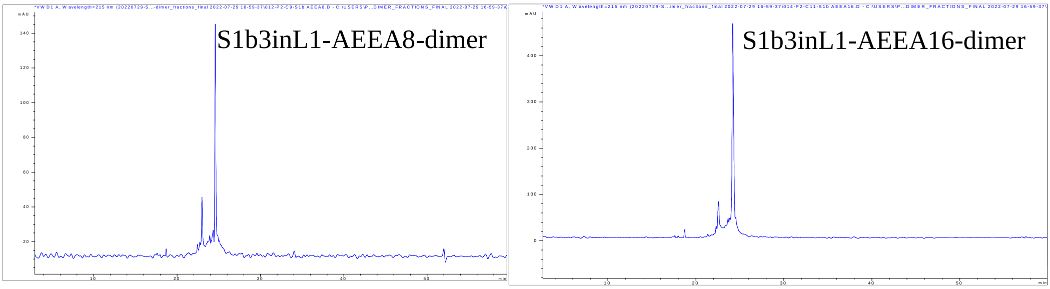

Figure S20. HPLC analysis of purified dimer peptides. UV-absorption traces at 215 nm. The main contaminant peak is an isobaric product assumed to be an alternate interstrand cyclisation.

### Tables S1-S5

Table S1. Overview of conditions used in the pre-clear and binding steps of the selection.

| Selection round | Pre-clear conditions | Binding conditions |
|---|---|---|
| R1 | - | 1 hour, 4°C |
| R2 | 3 x 20 min, RT | 1 hour, 4°C |
| R3 | 3 x 20 min, RT | 1 hour, 4°C |
| R4 | 3 x 20 min,RT | 1 hour, 4°C |
| R5 | 6 x 10 min, RT | 30 min, 4°C |
| R6 | 6 x 10 min, RT | 30 min, 4°C |
| Serum R1 | 3 x 20 min, RT | 30 min, 4°C (50% v/v Fetal calf serum) |
| Serum R2 | 3 x 20 min, RT | 30 min, 4°C (50% v/v Fetal calf serum) |
| RBD R1 (input R3) | 6 x 10 min, RT | 30 min, 4°C |
| RBD R2 (input R3) | 6 x 10 min, RT | 30 min, 4°C |
| RBD R1 (input R5) | 6 x 10 min, RT | 30 min, 4°C |
| RBD R2 (input R5) | 6 x 10 min, RT | 30 min, 4°C |
| SARS1 R1 | 6 x 10 min, RT | 30 min, 4°C |
| SARS1 R2 | 6 x 10 min, RT | 30 min, 4°C |
| SARS1 R3 (D-library) | 12 x 10 min, RT | 30 min, 4°C |

Table S2 List of synthesized peptides with the calculated and observed mass used in this study.

| **Peptide** | **Sequence** | **Calculated M+2H** | **Observed M+2H** |
|---|---|---|---|
| SARS2D1 | *Cyclo*(Ac-yNARILQRWFPELAHLC)G-$NH_2$ | 1113.6 | 1114.3 |
| SARS2D2 | *Cyclo*(Ac-yLC)LLFGRLHYHVHGPAG-$NH_2$ | 1046.5 | 1046.8 |
| SARS2D3 | *Cyclo*(Ac-yIC)TVRFWSPEWYPSFAG-$NH_2$ | 1124.5 | 1125.2 |
| SARS2D4 | *Cyclo*(Ac-ySIENYLKKLGYVLLWC)G-$NH_2$ | 1101.1 | 1101.4 |
| SARS2L1 | *Cyclo*(Ac-YIVWHDSLYSTHVQYVC)G-$NH_2$ | 1105.0 | 1105.3 |
| SARS2L2 | *Cyclo*(Ac-YVIWHPDRWTSVVLQIC)G-$NH_2$ | 1106.0 | 1106.4 |
| SARS2L3 | *Cyclo*(Ac-YSVVFSADGRHYYWDSC)G-$NH_2$ | 1076.0 | 1076.2 |
| SARS2R3L1 | *Cyclo*(Ac-YLVFHDNVYTVTVVHVC)G-$NH_2$ | 1052.5 | 1052.8 |
| SARS2R3L2 | *Cyclo*(Ac-YIVLHDGLRSATVLWIC)G-$NH_2$ | 1028.0 | 1028.3 |
| S1b D1 | *Cyclo*(Ac-yRC)LFGKVSWLGDYDDAG-$NH_2$ | 1052.5 | 1052.8 |
| S1b3inD1 | *Cyclo*(Ac-yLC)LFGKVSWCKPLDVCG-$NH_2$ | 1034.5 | 1034.8 |
| S1b3inD13 | *Cyclo*(Ac-yNC)LRGRVAWYSFRSEAG-$NH_2$ | 1087.5 | 1087.8 |
| S1b3inD22 | *Cyclo*(Ac-yTTVFC)RYFSAFARKSAG-$NH_2$ | 1057.5 | 1057.8 |
| S1b3inD31 | *Cyclo*(Ac-yYWYGC)RAHVYSWKLAAG-$NH_2$ | 1117.0 | 1117.4 |
| S1b3inL1 | *Cyclo*(Ac-YRRPREQIIIGSLWVFC)G-$NH_2$ | 1116.6 | 1116.9 |
| Alanine scan | | | |
| Pos2 | *Cyclo*(Ac-YARPREQIIIGSLWVFC)G-$NH_2$ | 1074.1 | 1074.4 |
| Pos3 | *Cyclo*(Ac-YRAPREQIIIGSLWVFC)G-$NH_2$ | 1074.1 | 1074.4 |
| Pos4 | *Cyclo*(Ac-YRRAREQIIIGSLWVFC)G-$NH_2$ | 1103.6 | 1104.2 |
| Pos5 | *Cyclo*(Ac-YRRPAEQIIIGSLWVFC)G-$NH_2$ | 1074.1 | 1074.8 |
| Pos6 | *Cyclo*(Ac-YRRPRAQIIIGSLWVFC)G-$NH_2$ | 1087.6 | 1088.3 |

| | | | |
|---|---|---|---|
| Pos7 | *Cyclo*(Ac-YRRPREAIIIGSLWVFC)G-$NH_2$ | 1088.1 | 1088.4 |
| Pos8 | *Cyclo*(Ac-YRRPREQAIIGSLWVFC)G-$NH_2$ | 1095.6 | 1096.2 |
| Pos9 | *Cyclo*(Ac-YRRPREQIAIGSLWVFC)G-$NH_2$ | 1095.6 | 1096.2 |
| Pos10 | *Cyclo*(Ac-YRRPREQIIAGSLWVFC)G-$NH_2$ | 1095.6 | 1096.3 |
| Pos11 | *Cyclo*(Ac-YRRPREQIIIASLWVFC)G-$NH_2$ | 1123.6 | 1124.3 |
| Pos12 | *Cyclo*(Ac-YRRPREQIIIGALWVFC)G-$NH_2$ | 1108.6 | 1108.9 |
| Pos13 | *Cyclo*(Ac-YRRPREQIIIGSAWVFC)G-$NH_2$ | 1095.6 | 1096.2 |
| Pos14 | *Cyclo*(Ac-YRRPREQIIIGSLAVFC)G-$NH_2$ | 1059.2 | 1059.4 |
| Pos15 | *Cyclo*(Ac-YRRPREQIIIGSLWAFC)G-$NH_2$ | 1102.6 | 1102.9 |
| Pos16 | *Cyclo*(Ac-YRRPREQIIIGSLWVAC)G-$NH_2$ | 1078.6 | 1079.3 |

Table S3 Change in melting temperature (ΔTm) measured by thermal shift analysis of indicated peptides (10 µM) against full length SARS-CoV-2 spike.

| Peptide | Sequence | ΔTm (°C) |
|---|---|---|
| SARS2L1 | *Cyclo*(Ac-YIVWHDSLYSTHVQYVC)G-$NH_2$ | 2 |
| SARS2L2 | *Cyclo*(Ac-YVIWHPDRWTSVVLQIC)G-$NH_2$ | 2 |
| SARS2D3 | *Cyclo*(Ac-yIC)TVRFWSPEWYPSFAG-$NH_2$ | 1 |
| SARS2D1 | *Cyclo*(Ac-yNARILQRWFPELAHLC)G-$NH_2$ | 0 |
| SARS2D2 | *Cyclo*(Ac-yLC)LLFGRLHYHVHGPAG-$NH_2$ | 0 |
| SARS2L3 | *Cyclo*(Ac-YSVVFSADGRHYYWDSC)G-$NH_2$ | 0 |
| S1b3inD1 | *Cyclo*(Ac-yLC)LFGKVSWCKPLDVCG-$NH_2$ | 0 |
| S1b3inD13 | *Cyclo*(Ac-yNC)LRGRVAWYSFRSEAG-$NH_2$ | 0 |
| S1b3inD22 | *Cyclo*(Ac-yTTVFC)RYFSAFARKSAG-$NH_2$ | 0 |
| S1b3inD31 | *Cyclo*(Ac-yYWYGC)RAHVYSWKLAAG-$NH_2$ | -0.5 |
| SARS2D4 | *Cyclo*(Ac-ySIENYLKKLGYVLLWC)G-$NH_2$ | -0.5 |

Table S4. Summary of cryo-EM data acquisition, image processing and model refinement statistics.

| | |
|---|---|
| **Data Collection** | |
| Microscope | Titan Krios G4 |
| Voltage (keV) | 300 |
| Nominal magnification | 130,000x |
| Movie acquisition rate | ~286 per hour |
| Detector | Falcon 4 |
| Energy filer | Selectris-X |
| Slit width (eV) | 10 |
| Calibrated pixel size (Å) | 0.929 |
| Cumulative exposure (e/Å$^2$) | 49 |
| Dose rate (e/pixel/sec) | 7.19 |
| Underfocus range (∝m) | 0.75-1.5 |
| Micrographs collected | 2612 |
| **Reconstruction** | |
| Final particles (no.) | 38,457 |
| Symmetry | C1 |
| B-factor (Å$^2$) | -55 |
| **Resolution (Å)** | |
| FSC 0.5 (masked) | 3.8 |
| FSC 0.143 (masked) | 3.2 |
| Resolution range (local) | 2.9-12.7 |
| **Refinement** | |
| Protein residues/atoms | 3246/25,232 |
| N-glycans/atoms | 49/686 |
| **Resolution (Å)** | |
| FSC 0.5 | 3.1 |
| FSC 0.143 | 2.9 |
| **Map correlation coefficient** | |
| Mask | 0.79 |
| Box | 0.76 |
| Volume | 0.77 |
| Peaks | 0.70 |
| **R.M.S. deviations** | |
| Bond Lengths (Å) | 0.004 |
| Bond Angles (°) | 0.490 |
| **MolProbity** | |
| Overall score | 1.46 |
| Clashscore | 5.31 |
| Ramachandran outliers (%) | 0 |
| Ramachandran favoured (%) | 96.97 |
| Rotamer outliers (%) | 0 |
| C-beta outliers | 0 |
| **EMRinger Score** | 2.77 |
| **Privateer** | |
| Wrong anomer | 0 |
| Wrong configuration | 0 |
| Unphysical puckering amplitude | 0 |
| In higher-energy conformations | 0 |

Table S5 Calculated and observed masses for the S1b3inL1 dimers

| AEEA repeats | Calc. $[M+8H]^{8+}$ | Obs. $[M+8H]^{8+}$ | Calc. $[M+7H]^{7+}$ | Obs. $[M+7H]^{7+}$ | Calc. $[M+6H]^{6+}$ | Obs. $[M+6H]^{6+}$ | Calc. $[M+5H]^{5+}$ | Obs. $[M+5H]^{5+}$ | Calc. $[M+4H]^{4+}$ | Obs. $[M+4H]^{4+}$ |
|---|---|---|---|---|---|---|---|---|---|---|
| 4 | | | 745.4 | 745.9 | 869.5 | 870.0 | 1043.1 | 1043.7 | 1303.7 | 1304.0 |
| 8 | 724.9 | 725.4 | 828.3 | 828.8 | 966.2 | 966.7 | 1159.2 | 1159.8 | 1448.8 | 1449.5 |
| 12 | 797.4 | 798.0 | 911.2 | 911.8 | 1062.9 | 1063.5 | 1275.3 | 1276.0 | 1593.8 | 1594.5 |
| 16 | 870.0 | 870.5 | 994.1 | 994.7 | 1159.6 | 1160.3 | 1391.3 | 1392.1 | 1738.9 | 1739.8 |

19. J. Jumper, R. Evans, A. Pritzel, T. Green, M. Figurnov, O. Ronneberger, K. Tunyasuvunakool, R. Bates, A. Žídek, A. Potapenko, A. Bridgland, C. Meyer, S. A. A. Kohl, A. J. Ballard, A. Cowie, B. Romera-Paredes, S. Nikolov, R. Jain, J. Adler, T. Back, S. Petersen, D. Reiman, E. Clancy, M. Zielinski, M. Steinegger, M. Pacholska, T. Berghammer, S. Bodenstein, D. Silver, O. Vinyals, A. W. Senior, K.

Kavukcuoglu, P. Kohli, D. Hassabis, Highly accurate protein structure prediction with AlphaFold. *Nature*. **596**, 583–589 (2021).
